# The protozoan commensal *Tritrichomonas musculis* is a natural adjuvant for mucosal IgA

**DOI:** 10.1101/2022.10.08.511442

**Authors:** Eric Yixiao Cao, Kyle Burrows, Pailin Chiaranunt, Ana Popovic, Xueyang Zhou, Cong Xie, Ayushi Thakur, Graham Britton, Matthew Spindler, Louis Ngai, Siu Ling Tai, Dragos Cristian Dasoveanu, Albert Nguyen, Jeremiah J. Faith, John Parkinson, Jennifer L. Gommerman, Arthur Mortha

## Abstract

Immunoglobulin(Ig) A antibodies are the most abundant antibodies supporting mucosal immune homeostasis and host-microbiota interactions. Driven by gut commensal microbes, IgA-secreting plasma cells (PC) differentiate through T cell-dependent (Td) or T cell independent (Ti) mechanisms. While commensal bacteria within the microbiota are known for their ability to promote IgA, the role of non-bacterial commensal microbes on the induction of IgA remains elusive. Here, we demonstrate that permanent colonization with the protozoan commensal *Tritrichomonas musculis* (*T.mu*) promotes T-cell dependent, IgA class-switch recombination and intestinal accumulation of IgA-secreting PC. *T.mu* colonization specifically drives the expansion of T follicular helper cells and a unique ICOS^+^ non-Tfh cell population, accompanied by an increase in germinal center B cells. Blockade of ICOS:ICOSL co-stimulation or MHCII-expression on B cells are central for the induction of IgA following colonization by *T.mu*, implicating a previously underappreciated mode of IgA induction following protozoan commensal colonization. Finally, the commensal *T.mu* further improves the induction of IgA-secreting plasma cells and their peripheral dissemination, even against non-protozoan, orally ingested antigens, identifying *T.mu* as natural adjuvant for IgA. Collectively, these findings propose a previously unknown, protozoa-driven mode of IgA induction that supports intestinal immune homeostasis even against non-microbial antigens.

## INTRODUCTION

The intestinal microbiota encompasses a diverse collection of microorganisms inclusive of viruses, bacteria, fungi, worms, and protozoa (1). While bacteria, fungi, worms and viruses have been recognized as regulators of the immune system, the contribution of commensal protozoan members of the microbiome remains less well understood (2). Recent reports characterized the murine commensal protozoa, *Tritrichomonas musculis (T.mu), Tritrichomonas muris (T.muris), Tritrichomonas grainger* (*T.grainger*) as agents of mucosal immune modulation (3–6). *Tritrichomonas* spp. have been shown to contribute to the regulation of the Th1 and Th17 response with beneficial and detrimental outcomes on the immunity and anti-microbial defense respectively (5, 7). Moreover, *Tritrichomonas* spp drive a tuft cell-ILC2 circuitry through microbial succinate to support anti-helminth immunity and epithelial regeneration (8). Surprisingly, colonization by *T.mu* was found to ameliorate central nervous inflammation in a mouse model of experimental autoimmune encephalomyelitis by promoting the accumulation of gut-derived plasma cells in the brain, collectively highlighting the multifactorial contributions of *Tritrichomonas* spp. in modulating host tissue and immune homeostasis (9).

Intestinal homeostasis requires interactions between the gut immune system and the commensal microbiota (10). IgA, the most prevalent isotype at mucosal surfaces, plays an essential role in maintaining the mucosal barrier, regulating bacterial growth, gene expression and host immunity to oral antigens (11). IgA exists in monomeric, dimeric and secreted forms, the latter, termed secretory IgA (SIgA). SIgA is comprised of an IgA-dimer, joined by a J-chain and the secretory component (SC), a polypeptide that stabilizes the Fc regions of IgA dimers. This complex forms following successful transportation of dimeric IgA across the intestinal epithelium via the polymeric-Ig receptor (PIGR)(12). Secreted into the lumen, SIgA binds to intestinal microbes and luminal antigens to mediate barrier support and homeostatic functions (13, 14). The intestinal lamina propria contains the largest population of IgA secreting PC, which develop through T cell-dependent (Td) and T cell-independent (Ti) class switch recombination (CSR). This process mediates differentiation of naïve B220^+^IgM^+^ B cells into IgA secreting PC in gut-associated lymphoid tissue (GALT), including *Peyer’s Patches* (PP), mesenteric lymph nodes (MLN) and isolated lymphoid follicles (ILF) (15, 16). During Ti IgA CSR, reactive oxygen species (ROS), retinoic acid (RA), *tumor necrosis factor alpha* (TNFα), *transforming growth factor beta* (TGFβ) interleukin (IL)-5, IL-6, IL-21, *tumor necrosis factor ligand superfamily members 13 (TNFSF13) and 13 b (TNFSF13B, BAFF)* increase chromatin accessibility of the IgA alpha locus in antigen-receptor stimulated B cells (17) In addition to antigen-receptor stimulation, Ti IgA CSR can be mediated via the stimulation of toll-like receptors (TLRs) on B cells (18). Conversely, Td IgA CSR requires the interaction of germinal center (GC) B cells with CD4^+^ T follicular helper (fh) cells that regulate the production of IgA through T cell antigen receptor MHCII interaction and costimulatory receptor-ligand pairs CD40-CD40L or ICOS-ICOSL (19). Td IgA CSR is believed to depend on antigen presentation by Dendritic cells (DC) and Follicular Dendritic Cells (FDC), providing a template for the selection of high-affinity GC B cells to generate antibody secreting plasma cells (20–22). Both pathways lead to the differentiation of GC B cells into PC and results in the migration of IgA secreting PC into the lamina propria. However, clear differences in affinity and reactivity against commensal microbes have been reported when comparing either Ti and Td IgA CSR in mice (23). This suggests that selective engagement of these pathways shape the reactivity of IgA towards commensal microbes. The generation of IgA within the gut is supported by the gut microbiota and readily able to adapt to new microbial members colonizing the intestinal tract (24). Interestingly, a once reactivity against gut bacteria has been established, the reactivity persists even if the colonization by the targeted microbe is transient (25). Moreover, IgA-specificity against intestinal bacteria has been shown to identify disease-promoting bacteria in Crohn’s disease patients supporting the notion that IgA serves as barrier sustaining agent for intestinal homeostasis (26).

However, the composition of the intestinal microbiota is not limited to bacteria, and the production and generation of IgA secreting PC driven by non-bacterial microorganism serves as a putative additional determinant of intestinal homeostasis (2). Gut protozoa like Tritrichomonads are an underappreciated component of the intestinal microbiota even in humans, working across many yet to be identified mechanisms that support the adaptive immune system including the activation of IgA secreting PC. We thus hypothesized that intestinal protozoa can regulate the mucosal IgA response.

In this study, we report that the murine protozoa *Tritrichomonas musculis* regulates the intestinal and systemic IgA responses. We characterize the previously unknown cellular and molecular mechanisms, and their implications in *T.mu*-driven IgA CSR, host-microbiome interactions and mucosal homeostasis. We demonstrate that colonization by *T.mu* increases luminal IgA levels and CD138^+^IgA^+^ PC counts, accompanied by higher numbers of PD-1^+^CXCR5^+^ Tfh cells and GL-7^+^FAS^+^ GC B cells in GALT. IgA induction following protozoan colonization requires Td IgA CSR in an ICOS:ICOSL and MHCII-TCR-dependent fashion. Subsequent analysis of the 16S rDNA gene sequences of IgA-coated gut bacteria indicates that *T.mu* colonization shifts the anti-bacterial IgA reactivity. Moreover, elevation of IgA after colonization with *T.mu* improved the induction and peripheral dissemination of IgA secreting PC specific to orally ingested food antigens, collectively demonstrating that *T.mu* colonization acts as a natural adjuvant for IgA by potentiating the systemic and mucosal IgA response.

## RESULTS

### Increased IgA levels following colonization with *T.mu*

Mice inoculated with the intestinal commensal protozoa *T.mu* show life-long colonization and do not develop signs of spontaneous inflammation and pathology (5). Previous reports demonstrated that colonization of the intestinal tract by newly colonizing gut bacteria impacts the mucosal IgA response (25). We therefore investigated if colonization by *T.mu*, as a non-bacterial commensal, could similarly change the mucosal IgA response. We first sought to determine serum IgA antibody levels of mice orally colonized by *T.mu* after 21 days, a timepoint that enables stable engraftment and sufficient time for the production of new antibodies to arise. ELISA for serum IgA revealed a significant increase in IgA levels after 3 weeks of colonization (Fig.1A). IgA-secreting plasma cells constitute a large fraction of lamina propria-resident immune cells in conventional mice. We next decided to determine whether colonization with *T.mu* would increase the quantities of IgA secreting cells in the intestinal lamina propria. Immunofluorescence imaging of IgA, followed by quantification of IgA^+^ cells indicated a significant increase in IgA-producing cells within the lamina propria of *T.mu* colonized mice (Fig.1B). Transcytosis of IgA into the gut lumen is critically dependent on the expression of the *polymeric Ig receptor* (PIGR) on intestinal epithelial cells (IEC) (12). As engraftment of *T.mu* into the intestinal microbiota increased serum IgA levels and IgA^+^ cells in the lamina propria, we hypothesized that luminal IgA levels and the expression of the IgA export machinery would be elevated in *T.mu* colonized mice. In support of this hypothesis, luminal IgA levels and the expression of *Pigr* showed a significant upregulation following colonization by *T.mu* (Fig.1C and D). These findings demonstrate that stable engraftment of *T.mu* into the intestinal tract promotes IgA production and transport across the intestinal epithelium. PC express the surface marker CD138 and are the major source of intestinal IgA (27, 28). To quantify absolute numbers of IgA^+^ PC, lamina propria leukocytes were isolated and IgA and CD138 co-expressing cells quantified. The lamina propria of *T.mu* colonized mice displayed a striking elevation in IgA^+^CD138^+^ PC compared to *T.mu*-free mice (Fig.1E). As IgA-secreting cells can be generated through Td and Ti pathways in gut-draining lymph nodes, we determined GC B cell and Tfh cell numbers in the mesenteric lymph nodes (MLN) of control and *T.mu*-colonized mice. In line with elevated IgA PC numbers, the MLN of *T.mu*-colonized mice showed higher proportions of FAS^+^GL-7^+^ GC B cells and PD-1^+^CXCR5^+^ Tfh cells indicating a Td contribution to the elevated IgA levels (Fig.1F and G). The increase in PC, GC B cells and Tfh cell frequencies were also reflected by higher absolute numbers of these cells (Fig.1H). Regulatory Foxp3^+^ T cells (Treg) have previously been associated with elevated IgA levels in mice, however, despite of higher IgA levels, colonization by *T.mu* failed to result in a significant increase in Tregs numbers in the lamina propria or MLN (Fig.S1A). The observation of unchanged numbers of Tregs were further supported by steady TFG-β levels in the intestinal tract, a factor known to promote Treg differentiation and IgA CSR (Fig.S1B)(29, 30). TNFSF13B and TNFSF13, two TNF superfamily members previously reported to promote Ti CSR to IgA, did not change in expression in the presence of *T.mu* (Fig.S1C). However, elevated levels of *Nos2* expression within the gut suggested a possible involvement of Ti CSR in the generation of IgA following colonization by *T.mu* (Fig.S1C)(31). We previously reported an increase in TNF production in *T.mu-*colonized mice, further supporting the idea of Ti CSR as mediator of the elevated IgA response (5, 7). Collectively, these results suggest both Td and Ti contribute to the increase in IgA observed in mice colonized with *T.mu*.

**Figure 1.**
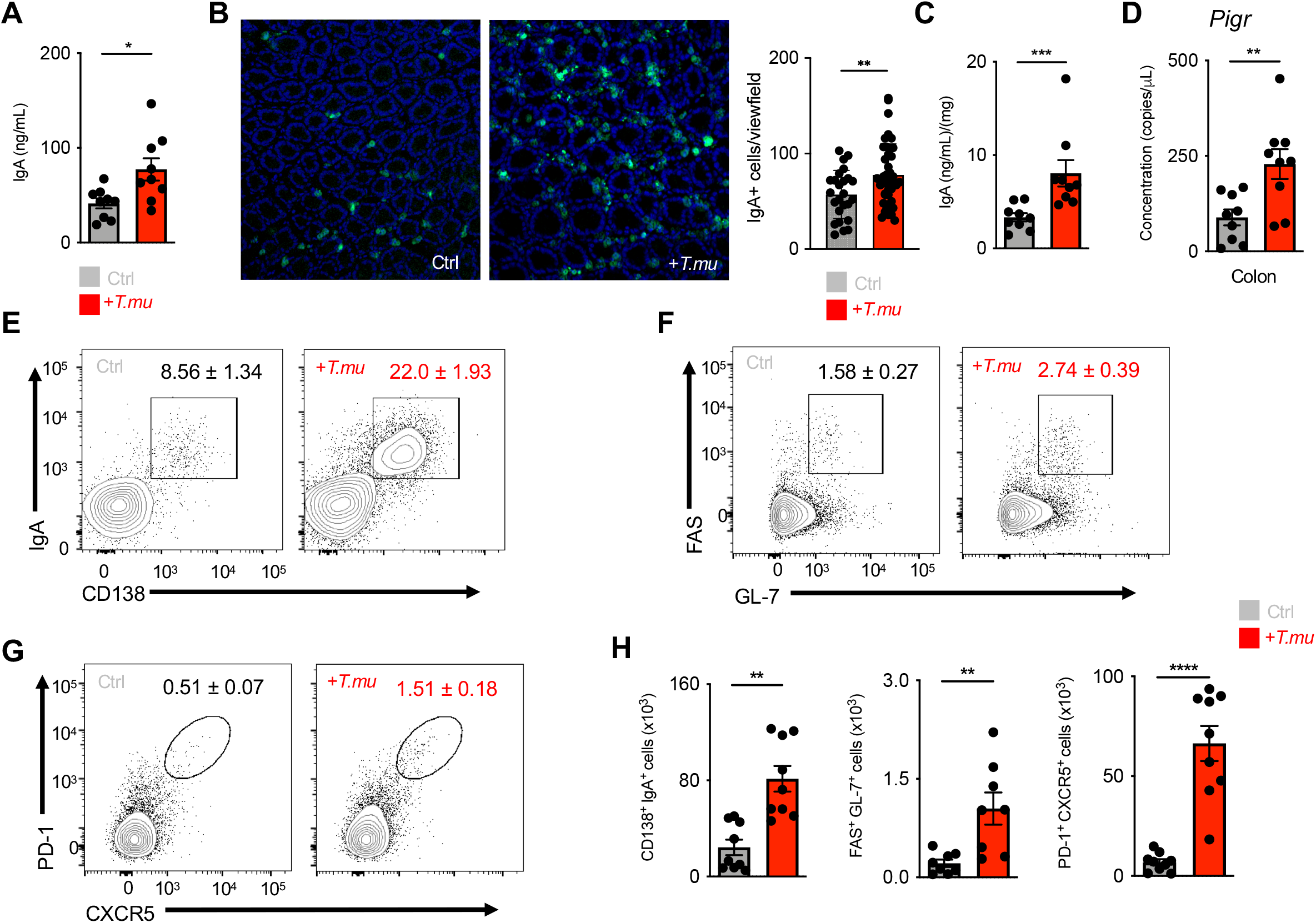
*Tritrichomonas musculis* induces mucosal IgA. **A)** Serum IgA levels in control and *T.mu*-colonized mice. **B)** Representative immunofluorescence images of colonic tissue sections from control (left) and *T.mu*-colonized mice (right) samples. Section were stained with anti-IgA antibodies. Adjacent bar graph shows quantification of IgA^+^ cells. **C)** IgA levels in fecal samples. **D**) Gene expression of *Pigr* in whole colonic tissue. **E)** Representative contour plots of lamina propria leukocytes stained with IgA and CD138. Gates identify IgA^+^CD138^+^ plasma cells. **F)** Contour plots show FAS^+^GL-7^+^ GC B cells in the MLN of control or *T.mu*-colonized mice. **G)** Representative flow cytometry plots identifying PD-1^+^CXCR5^+^ Tfh cells in the MLN of control or *T.mu*-colonized mice. Numbers adjacent to gates show percentages ±SEM in **E-G. H)** Quantification of plasma cells, GC B cells and Tfh cells. Data from at least 3 independent experiments with 3-4 mice/group are shown. Mann-Whitney U test was performed. *= p ≤ 0.05, **= p < 0.01, ***=p < 0.001, **** = p < 0.0001, NS = not significant.

### *T.mu*-driven IgA is ICOS and T cell-dependent

To dissect the contribution of Td and Ti IgA CSR to the increase in IgA following colonization by *T.mu*, we first colonized *Nos2*^*−/−*^ or *Tnfa*^*−/−*^ mice and their respective age-matched, heterozygous littermate controls with *T.mu* to assess their lamina propria IgA PC numbers. Both *Nos2* and *Tnfa* were dispensable for the *T.mu*-driven induction of IgA, as both *Nos2*^*−/−*^ or *Tnfa*^*−/−*^ and their heterozygous littermate controls showed an equal induction of IgA after *T.mu* engraftment into the microbiota. The findings prompted us to investigate whether genetic abrogation of T cell development, or antibody-mediated depletion of CD4^+^ T cells would prevent the increased IgA levels following colonization with *T.mu* (Fig.2A and B). To address this, groups of C57Bl/6 and *Tcrb*^*−/−*^ were either left untreated or colonized with *T.mu* for 3 weeks. Additional groups of C57Bl/6 mice received control Ig or depleting anti-CD4 Ig injections for the course of 3 weeks starting 3 day prior to *T.mu* colonization. Measuring serum and fecal IgA levels in T cell-deficient or T cell-depleted mice revealed a failure to increase IgA levels, specifically in groups colonized with *T.mu* (Fig.2C and D). To causally demonstrate the requirement of CD4 T helper cells in the *T.mu* driven IgA induction, naïve, splenic, FACS-purified CD4^+^ T cells from wild type mice were injected into *Tcrb*^*−/−*^ control mice, or *Tcrb*^*−/−*^ mice colonized with *T.mu*. Analysis of IgA^+^ PC frequencies and numbers demonstrated the requirement of CD4+ T cells and Td IgA CSR in promoting IgA production following colonization with *T.mu* (Fig.2E).

**Figure 2.**
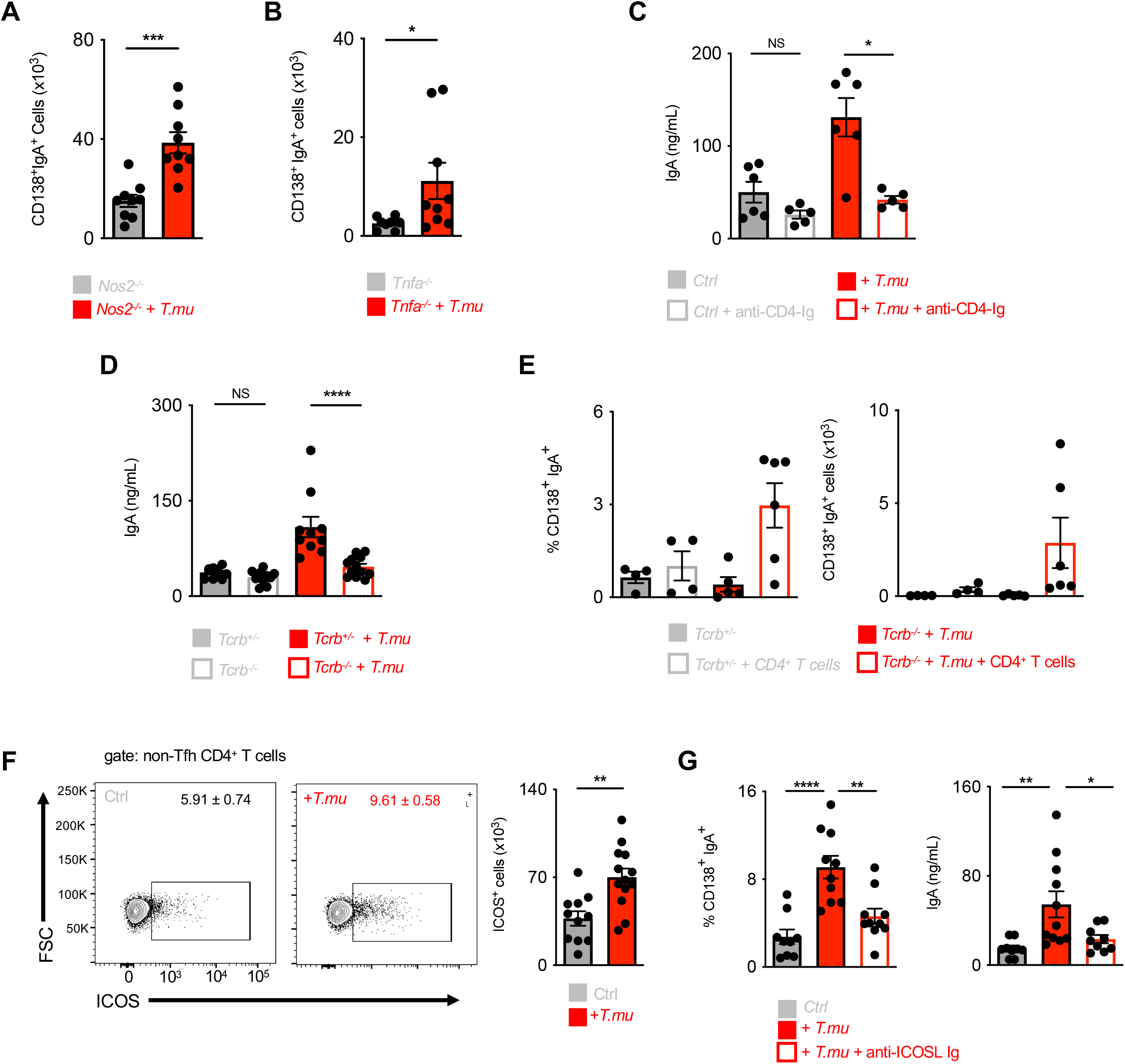
*T.mu* induces IgA in an ICOS and T cell-dependent fashion. **A)** Lamina propria IgA^+^CD138^+^ plasma cell numbers in control *Nos2*^*−/−*^ mice or *Nos2*^*−/−*^ mice colonized with *T.mu*. **B)** Lamina propria IgA^+^CD138^+^ plasma cell numbers in control *Tnfa*^*−/−*^ mice or *Tnfa*^*−/−*^ mice colonized with *T.mu*. **C)** Serum and fecal IgA levels in control mice or mice colonized with *T.mu*. groups of mice were injected with depleting anti-CD4-Ig. **D)** Serum and fecal IgA levels in *Tcrb*^*−/−*^ control mice or *Tcrb*^*−/−*^ mice colonized with *T.mu*. **E)** Quantification of lamina propria IgA^+^CD138^+^ plasma cells in *Tcrb*^*−/−*^ control mice or *Tcrb*^*−/−*^mice colonized with *T.mu*. **F)** Contour plots show ICOS expression on non-Tfh cells in MLN from control mice or mice colonized with *T.mu*. Bar graph adjacent shows quantification of absolute ICOS^+^ non-Tfh cells. **G)** Quantification of IgA^+^CD138^+^ plasma cells and serum IgA levels in mice control mice, mice colonized with *T.mu* and *T.mu*-colonized mice injected with blocking anti-ICOSL-Ig. Data shown is from 2-3 independent experiments with 3-4 mice/group. Mann-Whitney U test was performed. *= p ≤ 0.05, **= p < 0.01, ***=p < 0.001, **** = p < 0.0001, NS = not significant.

We next determined if the increase in GALT-resident Tfh cells may be accompanied by changes in the expression of IgA CSR-promoting factors. To test this, we sorted Tfh cells from control mice and *T.mu* colonized mice and determined CD40L expression (Fig.S1D), *Il4, Il6, Il13, Il21, Il10* expression (Fig.S1E), or *Tbx21* and *Gata3* expression (Fig.S1F). Surprisingly, no changes were recorded indicating an unaltered Tfh cell phenotype in the presence of *T.mu*. CSR has been reported to also occur in extrafollicular regions of the lymph nodes and spleen, where ICOS-expressing non-Tfh cells are suspected to promote this process (32, 33). Supporting the idea that ICOS-expressing cells promote IgA, a recent report demonstrated that ICOSL deficiency minimizes IgA levels and anti-microbial IgA recognition (34). We therefore assessed ICOS expression on non-Tfh CD4^+^ T cells and observed an increase in ICOS-expressing CD4^+^ non-Tfh cells in the presence of *T.mu* supporting the hypothesis that *T.mu*-driven IgA requires ICOS:ICOSL interactions. (Fig.2F). To determine if ICOS expression on non-Tfh cells is a consequence of microbial colonization, we analyzed the CD4^+^ T cell compartment of germ-free mice following colonization with previously reported IgA inducing commensal bacteria. We colonized germ-free mice with six distinct *Bacteroides ovatus* strains (A, C, H, I, L, N), that covered a graded capability to promote IgA CSR (A > C > H > I > L > N)(35). Three weeks post colonization MLN PD-1 ^+^CXCR5^+^ Tfh cells and ICOS^+^ non-Tfh cells were quantified. *B.ovatus* strains previously reported to induce high levels of IgA failed to display an increase in Tfh cells and ICOS-expressing Tfh cells that would be comparable to mice colonized by *T.mu* (Fig.S1G and H). These results indicate a unique mode of immune regulation following *T.mu* colonization that drives IgA in the presence of commensal protozoa. To investigate the importance of ICOS on the induction of IgA following colonization by *T.mu*, we abrogated ICOS:ICOSL interactions through injections of blocking anti-ICOSL antibodies. Strikingly, this treatment almost completely reversed the elevated serum IgA levels and lamina propria IgA plasma cells numbers in mice colonized with *T.mu* (Fig.2G). In conclusion, these results demonstrate that *T.mu*-mediated IgA induction operates through ICOS-expressing Tfh and non-Tfh cells. A reduced anti-bacterial IgA reactivity, as a consequence of a defective ICOS:ICOSL interaction has previously been reported, suggesting that colonization by the protozoan commensal *T.mu* may improve anti-bacterial IgA recognition through the upregulation of ICOS on CD4^+^ T cells and the increase in Tfh cells.

### Shift in microbial composition and IgA-reactome following colonization with *T.mu*

To test the hypothesis that colonization with *T.mu* would change the anti-bacterial IgA reactivity, fecal bacteria were isolated from control or *T.mu* colonized mice to stain their intestinal bacteria with anti-IgA antibodies to determine the degree of IgA coating. Mice colonized by *T.mu* displayed significantly higher IgA coating compared to control mice, supporting Landuyt *et al*.’s data of an ICOS:ICOSL-dependent regulation of anti-microbial IgA (34)(Fig.3A). Sera obtained from control mice and mice colonized with *T.mu* recapitulated the elevated IgA-coating pattern even when fecal bacteria from antibody-deficient *Rag2*^*−/−*^ mice were used as template for IgA coating (Fig.S2A). These findings demonstrate that serum IgA and secreted IgA display an elevated capacity to bind bacteria after colonization with *T.mu*. To examine whether anti-commensal IgA antibodies show a distinct reactivity towards luminal bacteria in the presence of *T.mu*, equal concentrations of fecal protein from bacterial lysates of conventional C57Bl/6 mice were separated by SDS-PAGE and transferred onto a nitrocellulose membrane. Stripes of these membranes were separated and incubated with control serum, or serum collected from *T.mu* colonized mice. Anti-IgA binding to the blotted bacterial proteins was detected using anti-mouse IgA secondary antibodies. Membranes developed under identical conditions revealed that IgA antibodies from *T.mu* colonized mice exhibit a distinct band pattern when compared to IgA antibodies from control mice, showing that serum IgA antibodies from *T.mu* colonized mice have a distinct IgA reactivity against bacterial proteins (Fig.3B). These results inspired us to purify fecal IgA-coated and non-coated bacteria from control and *T.mu* colonized mice to determine their composition. We isolated DNA from FACS-sorted IgA-coated and non-coated bacteria and performed 16S rDNA gene sequencing on the respective libraries. Principal coordinate analysis, based on Bray-Curtis compositional dissimilarities, demonstrated separation of IgA-coated and non-coated bacteria in control and *T.mu* colonized mice (*p*<0.001). Bacterial composition obtained from mice colonized with *T.mu* clustered distinctly from control samples, supporting a change in the profile of IgA-reactive bacteria in the presence of the protist (Fig.3C). At week 3, although fecal bacterial richness was similar in naïve and *Tmu* colonized mice, both uncoated and more interestingly IgA-coated bacteria were significantly more diverse in *T.mu* colonized mice, suggesting a broader IgA reactivity in the presence of *T.mu*. Evenness displayed similar trends, albeit with elevated fecal diversity in control mice. While the bacterial evenness remained consistently distributed, increased richness of OTUs were observed in mice colonized by *T.mu* (Fig.S2B and 3D). These changes in richness were also reflected when comparing the relative abundance of taxa across the different experimental groups and specifically emphasize an expansion in the reactivity of IgA in the presence of *T.mu* (Fig.3D-E). An increase in Clostridia and Gammaproteobacteria were apparent within the fraction of IgA-coated bacteria when mice were colonized with *T.mu* (Fig.3E). These findings were recapitulated when comparing OTUs from the IgA-coated bacterial fraction, indicating that IgA-coating, driven by protozoan colonization, promotes a shift in the bacterial microbiota (Fig.S2C). Collectively, these findings show that colonization with the protozoan commensal *T.mu* changes the IgA reactivity and induces a shift in the abundance intestinal bacterial microbes.

**Figure 3.**
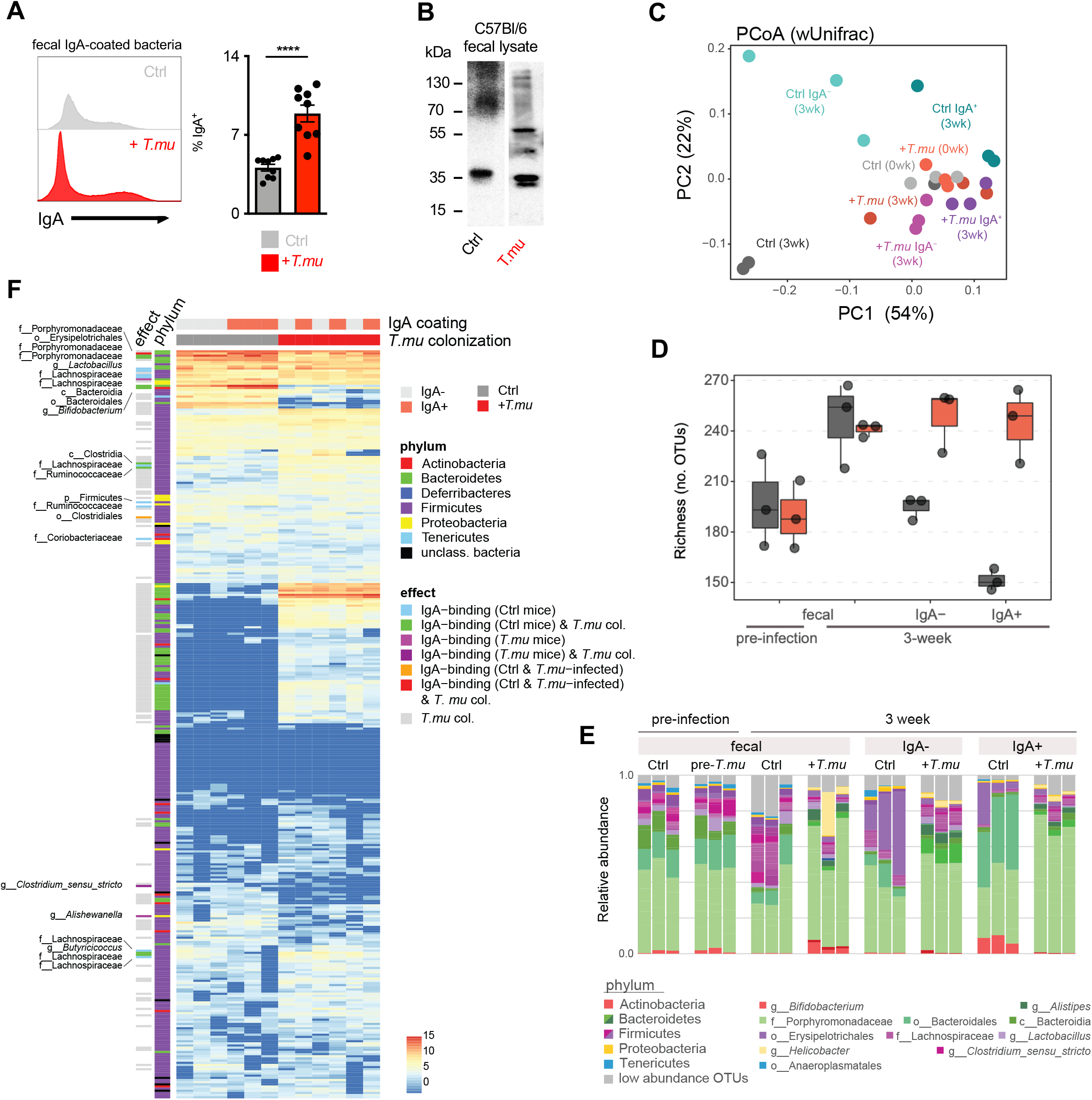
*T.mu* shapes the anti-bacterial IgA reactome. **A)** Fecal bacteria were isolated from control or *T.mu*-colonized mice and stained with anti-IgA antibodies. Histograms show staining of anti-IgA antibodies. Adjacent bar graph shows quantification of IgA-coated bacteria. **B)** Western blot analysis of IgA reactivity on wild type C57Bl/6 fecal bacterial protein lysates. Membranes were developed after incubation with serum from control mice or *T.mu*-colonized mice. Numbers adjacent to membranes indicate molecular weight. **C)** PCoA of 16S rDNA gene sequencing data obtained from control and *T.mu*-colonized samples. OTUs, IgA^−^ OTUs and IgA^+^ OTUs from control and *T.mu*-colonized mice were used **C-F)** Compositional dissimilarities of fecal and flow-sorted bacterial communities, based on weighted Unifrac distances. Analysis was carried out using samples rarefied to the lowest sample depth. **D)** Bacterial richness of fecal and flow-sorted microbiota. Values represent averaged diversities from 100 datasets rarefied to the minimum sample depth (21,000 reads). **E)** Relative abundance of bacterial OTUs, coloured by phylum as indicated. Select highly abundant taxa are indicated. Low abundance taxa (<1% relative abundance) were combined and are represented in grey. **F)** Normalized abundance of IgA-coated and uncoated taxa. Bacteria differentially abundant based on IgA coating or presence of T.mu in samples were determined using DESeq2, and significant effects indicated by coloured bars on the left, p<0.05. Bacterial phyla are and taxonomies of those significantly associated with IgA-coating are labelled, and taxon phyla and relevant effect are indicated in the colour key to the right.

### B cell-expressed MHCII is essential for *T.mu* driven IgA

Bacterial antigens are translocated through epithelial microfold cells and subsequently transferred to MHCII-expressing dendritic cells (DC) to facilitate the transport of the antigen to the T cell zone of the draining lymph node (14, 36). As ICOS expression on T cells and the expansion of Tfh cells have been demonstrated to require T cell receptor (TCR) engagement through MHCII and *T.mu* colonization elevates the numbers of ICOS^+^ CD4^+^ non-Tfh cells and Tfh cells, we reasoned that *T.mu*-mediated induction of IgA and ICOS may also depend on MHCII-mediated antigen presentation (37, 38).

To address this question, we left *MHCII*^*−/−*^ mice untreated, or colonized them with *T.mu* to determine ICOS^+^ non-Tfh cells, Tfh cell numbers and IgA levels. In contrast to age-matched littermate control mice, *MHCII*^*−/−*^ mice failed to show an increase in IgA, Tfh cells and ICOS-expressing non-Tfh cells (Fig.4A). These findings indicate that protozoa-driven IgA induction and T cell expansion require MHCII-TCR interactions. Naive B cells are underappreciated antigen-presenting cells and recently gained attention as a critical source of MHCII that promotes the anti-plasmodium antibody response (39). We hypothesized that B cell-presented antigens are mandatory for the ICOS-dependent, anti-commensal IgA response driven by colonization with *T.mu*. To test this hypothesis, we colonized B cell-deficient *μMT*^*−/−*^ mice with *T.mu* and determined the numbers of ICOS^+^ non-Tfh cells in the draining lymph nodes. In support of our hypothesis, we observed that *μMT*^*−/−*^ mice failed to show a significant increase in ICOS^+^ non-Tfh cells (Fig.4B). Tfh cells further required B cells for their expansion and consequently failed to increase in *μMT*^*−/−*^ mice after protozoan colonization, demonstrating that B cells are a crucial element in supporting the *T.mu*-driven expansion of MHCII-dependent, ICOS^+^ non-Tfh cells and Tfh cells (Fig.S3A). To determine whether MHCII on other hematopoietic cells (e.g. macrophages or DC) could contribute to the anti-commensal IgA response after colonization with *T.mu*, bone marrow chimeric mice were generated. A group of lethally irradiated *μMT*^*−/−*^ mice received a 50:50 mixture of *MHCII*^*−/−*^:*μMT*^*−/−*^ bone marrow cells while the control group of *μMT*^*−/−*^ mice was transplanted with a 50:50 mixture of *MHCII*^*+/−*^:*μMT*^*+/−*^ bone marrow cells. We hypothesized that a reconstituted *MHCII*^*−/−*^:*μMT*^*−/−*^ immune system, comprised of only MHCII-deficient B cells would fail to show an increase in the *T.mu*-driven IgA response. Following full bone marrow reconstitution, mice were orally colonized with *T.mu* and analyzed for mucosal IgA-producing plasma cells, GC B cells, Tfh cells and ICOS^+^ non-Tfh cells. In line with our hypothesis, and supported by the observations in *μMT*^*−/−*^ and *MHCII*^*−/−*^ mice, colonization by *T.mu* only yielded a full IgA response in mice capable of presenting antigen via B cell-specific MHCII (Fig.4C). These findings demonstrate that protozoan colonization initiates an ICOS- and T cell-dependent IgA response through MHCII antigen-presentation by B cells. Colonization with *T.mu* thus elevates GC B cells, Tfh cells and IgA PC, to promote new IgA reactivity against bacterial antigens. These observations inspired us to ask whether the underlying immune activation following colonization by *T.mu* could be harnessed to improve the IgA response against non-bacterial antigens in the form of a natural adjuvant.

**Figure 4.**
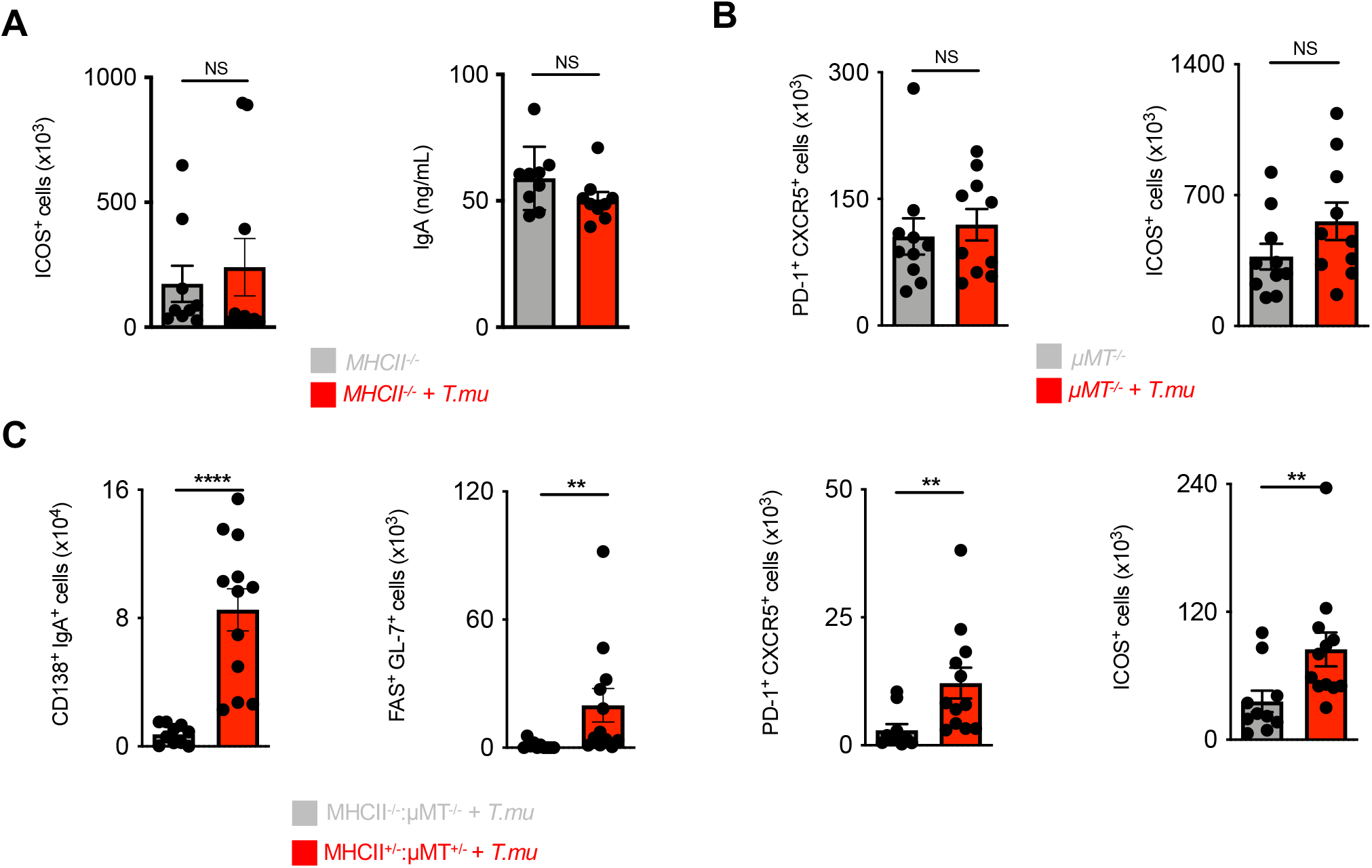
*T.mu*-driven IgA requires antigen-presentation by B cells. **A)** Quantification of ICOS^+^ non-Tfh cells in the MLN and serum IgA levels in control and *T.mu*-colonized *MHCII*^*−/−*^ mice. **B)** Quantification of MLN Tfh and ICOS^+^ non-Tfh cell numbers in *μMT*^*−/−*^ control mice or *μMT*^*−/−*^ mice colonized with *T.mu*. **C)** Analysis of bone marrow chimeric mice. Groups of mice received a 50:50 mixture of either *MHCII*^*−/−*^ *μMT*^*−/−*^ or *MHCII*^*+/−*^ *μMT*^*+/−*^ bone marrow cells and were colonized with *T.mu* after full bone marrow reconstitution. Absolute numbers of colonic IgA^+^CD138^+^ plasma cells, MLN FAS^+^GL-7^+^ GC B cells, PD-1^+^CXCR5^+^ Tfh cells and ICOS+ non-Tfh cells were quantified. Data shown is representative of at least 3 independent experiments with 3-4 mice/group. Mann-Whitney U test was performed. *= p ≤ 0.05, **= p < 0.01, ***=p < 0.001, **** = p < 0.0001, NS = not significant.

### Colonization with *T.mu* promotes peripheral dissemination of gut-primed plasma cells

Plasma cells, primed against gut luminal antigens, have been reported to egress from the lamina propria and migrate to peripheral tissues (9, 40). We reasoned that the increased anti-commensal IgA response driven by *T.mu* would also improve an IgA response against non-bacterial orally ingested antigens and mediate the peripheral dissemination of antigen-specific PC. To test our hypothesis, groups of mice were either left untreated or colonized with *T.mu* followed by oral exposure to the model antigen Ovalbumin (OVA). OVA-containing drinking water was provided *ad libidum* for 3 consecutive weeks followed by the analysis of OVA-specific serum IgA levels and OVA-specific IgA-secreting PC in peripheral organs. Oral exposure to OVA commonly elicits very weak antigen-specific IgA titers in conventional wild type mice, indicating that in the absence of protozoa, the microbiota has a weak adjuvant activity towards IgA (41). In line with these observations, mice free of the protozoan commensal *T.mu* displayed low levels of OVA-specific serum and fecal IgA antibodies (Fig.5A). However, mice colonized with *T.mu* showed significantly higher anti-OVA IgA levels in their serum and feces, demonstrating that the *T*.mu-boosted mucosal IgA response potentiates IgA against non-microbial, luminal antigens (Fig.5A). As anticipated, OVA treatment alone had no effects on total IgA levels (Fig.S4A). To investigate if anti-OVA IgA secreting PC would display an improved peripheral dissemination, ELISPOT assays of OVA-specific IgA PC were performed on MLN, spleen and bone marrow cells. In line with an increased serum and fecal anti-OVA IgA response, *T.mu* colonization further resulted in a significantly higher peripheral dissemination of OVA-specific IgA-secreting PC (Fig.5B). These findings demonstrate that the commensal protozoa *T.mu* functions as natural adjuvant for luminal antigens by boosting the TD induction of mucosal IgA and by enhancing the peripheral dissemination of gut-primed antigen-specific PC.

**Figure 5.**
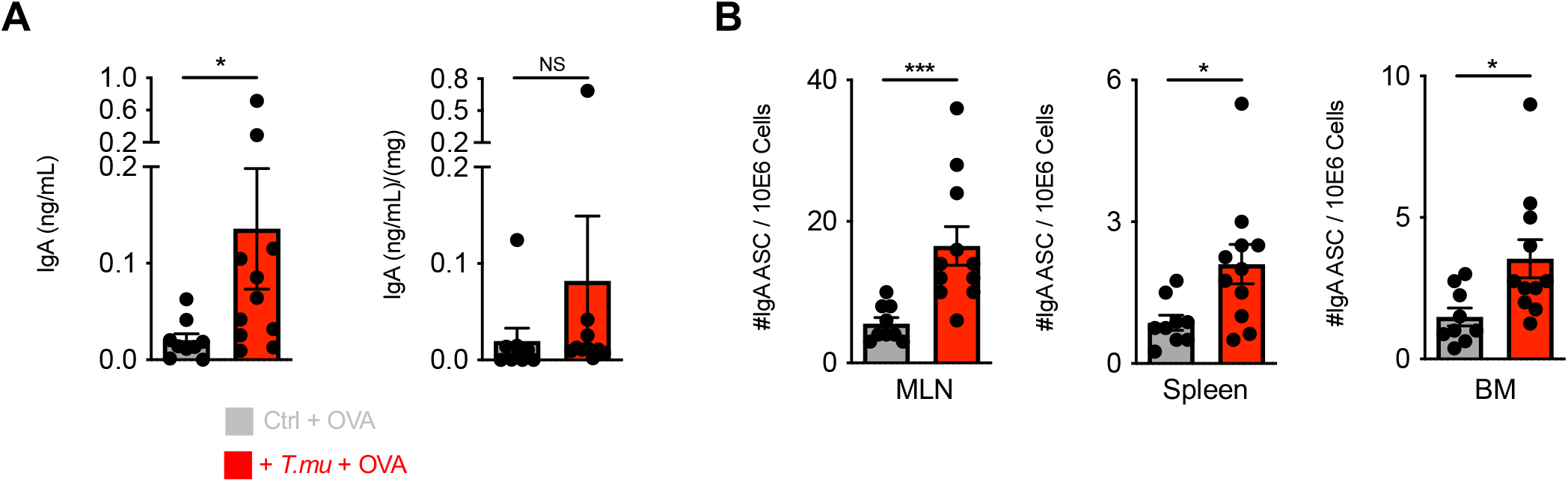
*T.mu* promotes peripheral dissemination of IgA-secreting plasma cells primed against gut antigens. **A)** Groups of control or *T.mu*-colonized mice were treated with OVA-containing drinking water ad libidum for 3 weeks. OVA-specific IgA levels were determined in the serum or feces. **B)** MLN, spleens and bone marrow were harvest from mice used in A) and numbers of OVA-specific anti-IgA secreting cells determined. Data shown is representative of at least 3 independent experiments with 3-4 mice/group. Mann-Whitney U test was performed. *= p ≤ 0.05, **= p < 0.01, ***=p < 0.001, **** = p < 0.0001, NS = not significant.

## DISCUSSION

Our study uncovers an underappreciate contribution by the recently identified protozoan commensal *T.mu* to the mucosal and systemic IgA. Even though the impact of the gut microbiota on the induction of IgA is appreciated, the impact of protozoan commensals within the microbiota has only been addressed by a limited number of studies, preferably focusing on pathogenic protozoa (42, 43). Here, we report that colonization with the commensal protozoa *T.mu* elicits substantial changes in the host’s IgA production, IgA reactivity and PC dissemination. The obtained elevations in IgA levels and IgA reactivity are the result of newly generated IgA-producing PC, arising through a Td mechanism dependent on ICOS:ICOSL co-stimulation, independent of IgA promoting bacteria. We demonstrate that serum and fecal IgA show greater coating of intestinal bacteria after protozoan colonization further indicating that *T.mu* colonization shapes the microbiota towards greater richness and evenness, and associates with a broader IgA reactivity towards intestinal commensal. The elevated production of IgA required B cell-expressed MHCII, suggesting B cells as critical antigen-presenting cell for the induction of IgA after colonization with *T.mu*. Importantly, the elevated induction of IgA in mice carrying *T.mu* additionally promotes IgA reactivity to non-bacterial, orally ingested antigens, accompanied by increased peripheral dissemination of PC primed against luminal antigens. These findings extend our understanding of how the microbiota controls the host’s adaptive immune response and adds non-bacterial commensal as important regulator of the homeostatic mucosal IgA response.

The role of *Tritrichomonas* spp. as driver of the host’s immune response has multiple effects on the intestinal tract. Colonization of mice with *T.mu* potentiates the anti-microbial immunity against the intracellular pathogen Salmonella (5, 7). However, the elevated threshold of immunity imposed by *T.mu* confers risks to the host’s health and can exacerbate disease outcomes under permissive circumstances (5, 6). In contrast to these findings is *T.mu*’s ability to promote the anti-inflammatory effects of gut-derived PC in the central nervous system (CNS) suggesting a divergent role of *T.mu* in mediating peripheral disease tolerance (9). While IgA is dispensable for the ameliorated pathology in the CNS, peripheral dissemination of PC and the release of other PC-related factors arise as intriguing targets for future research (9). Moreover, the expansion of ICOS-expressing non-Tfh cells warrants attention and suggests, either an increase in undifferentiated Tfh cells in the GALT of mice, or an independently arising T cell subset that promotes Td IgA CSR in an ICOS-dependent fashion (44). New experimental approaches are required to selectively address these observations and dissect the distinct contribution and developmental trajectories of ICOS-expressing non-Tfh cells. The observation that B cell-expressed MHCII contributes to the expansion of IgA PC and the germinal center response in the presence of *T.mu* is another exciting observation that implicates gut luminal antigens as driver of an increased IgA response through antigen presentation by B cells. While the role of B cells as inducer of IgA has been reported for a conserved epitope in the N-terminus of the circumsporozoite protein in *Plasmodium falciparum*, other protozoan or bacteria epitopes may also be the target of such response (39). These observations challenge the role of conventional Dendritic cells and follicular Dendritic cells as central antigen presenting cell for mucosal IgA responses and demand future investigations dissecting the distinct roles of each antigen presenting cell type to mucosal IgA production (31).

IgA responses against mucosal antigens are of growing interest for the development of vaccines against emerging mucosal pathogens (45). Considering that the microbiota impacts the systemic vaccine response, a stratification around a protozoan prebiotic formulation may be an attractive approach to tailor the outcome of vaccinations towards a sustained and improved IgA response. In line with this notion, supplementing the gut microbial community with a protozoan commensal like *T.mu* may serve as natural adjuvants for mucosal IgA and offer a promising approach to drive IgA responses to orally ingested antigens.

Collectively, we report multiple novel observations centered around the host’s IgA response in the presence of a previously underappreciated gut commensal microbe. The uncovered modes of action for a protozoan-boosted IgA response uses a Td pathway along the axis of ICOS:ICOSL co-stimulation and B-cell dependent MHCII expression. This interaction comes with the benefits to additionally promote IgA responses against non-microbial luminal antigen and may be of benefit to the host in tolerating new dietary antigens. Protozoan commensals like *T.mu*, thus are an interesting unexplored group of microbes to investigate innovative natural adjuvant formulations for the induction of IgA with beneficial effects for the induction of oral tolerance and host defense.

## ACKNOWLEDGEMENTS

We thank all members of the #onlylabever for their continued support and scientific insight. We would like to thank all members of the University of Toronto Flow Cytometry facility for their outstanding support and assistance, and thank the Centre for Analysis of Genome Evolution & Function (CAGEF), for DNA sequencing. We wish to thank the staff and resources of the Microbiome Translational Center, Icahn School of Medicine at Mount Sinai for exceptional support. The graphical abstract has been created using BioRender. This study was supported grants from the Canadian Foundation for Innovation John R. Evans Leaders Fund (A.M.), the Canadian Institutes of Health Research (CIHR; PJT-388337 to A.M., MRT-168043 to J.P. and A.M. and PJT-152921 to J.P.), the Natural Sciences and Engineering Research Council of Canada (NSERC; RGPIN-2019-04521 (A.M.) and RGPIN-2019-06852 to J.P.). Further funding was provided by an Ontario Trillium Scholarship and Vanier Canada Graduate Scholarship - NSERC to P.C.. K.B. is supported by a Canadian Institutes of Health Research (CIHR) Banting Postdoctoral Fellowship Program and a Canadian Allergy Asthma and Immunology Foundation Research Fellowship in Type 2 Inflammation supported by Sanofi Canada. L.N. is supported by an Ontario Graduate Scholarship and a NSERC-PGS award. S.L.T. is a recipient of the Dr. Edward Ketchum Graduate Student Scholarship and the Canada Graduate Scholarships – Master’s (CGS M) award. A.M. is the Tier 2 Canadian Research Chair in Mucosal Immunology and supported by the Tier 2 CRC-CIHR program (CRC-2021-00511).

## AUTHOR CONTRIBUTIONS

E.Y.C. wrote the manuscript, performed experiments, and analyzed the data. K.B., P.C., X.Z., C.X., A.T., G.B., M.S., L.N., S.L.T., N.A., D.C.D. and A.N. performed experiments. A.P. performed computational analysis of 16S rDNA gene sequencing data. J.J.F. provided critical reagents and experimental tools. J.P. supported bioinformatic analysis and funding acquisition. J.L.G. co-supervised this study. A.M. developed the concept of this study, funded the project, helped analyzing data, supervised the project, and wrote the manuscript.

## Supplementary Figure Legends

**Supplementary Figure 1. Unaltered Tfh cell phenotype in the presence of *T.mu***

**A)** Quantification of FOXP3 expression in CD4^+^ T cells in the lamina propria and MLN of control and *T.mu*-colonized mice. **B)** Quantification of biological active TGF-β in colonic tissue explants from control and *T.mu*-colonized mice. **C)** Gene expression analysis of *Tnfsf13, Tnfsf13b* and *Nos2* in total colonic tissue. **D)** CD40L expression on MLN PD-1 ^+^CXCR5^+^ Tfh cells from control or *T.mu*-colonized mice. **E)** PD-1 ^+^CXCR5^+^ Tfh cells were sorted form the MLN of control or *T.mu*-colonized mice. Gene expression analysis was performed for cytokines *Il4, Il6, Il13, Il21, Il10* and **F)** transcription factors *Gata3* and *Tbx21*. **G and H)** Groups of germ-free mice were colonized with *Bacteriodes ovatus* strains (A, H, N, I, C and L). Three weeks after colonization MLN PD-1 ^+^CXCR5^+^ Tfh cells (G) and ICOS^+^ non-Tfh cells (H) were quantified. Data shown is representative of at least 2 independent experiments.

**Supplementary Figure 2. Shift in anti-bacterial IgA reactivity following colonization with *T.mu***.

**A)** Bacterial evenness of fecal and flow-sorted microbiota. Values represent averaged diversities from 100 datasets rarefied to the minimum sample depth (21,000 reads). **B)** Serum from control mice or *T.mu*-colonized mice was collected and used to stain fecal bacteria obtained from *Rag2*^*−/−*^ mice. Histogram shows coating of fecal bacteria with serum IgA. Adjacent bar graphs show quantification of IgA-coated bacteria and the serum staining intensity displayed as mean fluorescent intensity normalized to controls. **C)** Heat map indicates microbial relative abundance pre and post sorted samples. Bars adjacent to heat map indicate effect-driving variables. Data shown is representative of at least 3 independent experiments.

**Supplementary Figure 3. B cell-dependent Tfh response in *T.mu*-colonized mice**.

**A)** Groups of *μMT*^*−/−*^ mice were colonized with *T.mu* or left untreated. 3 weeks post colonization, PD-1 ^+^CXCR5^+^ Tfh cells were quantified in the MLN. Data shown is representative of at least 3 independent experiments with n=4 mice/group.

**Supplementary Figure 4. Exposure to oral Ovalbumine does not alter *T.mu*-driven IgA levels**.

**A)** Total serum and fecal IgA levels were determined in groups of mice exposed to OVA-containing drinking water in the presence of absence of *T.mu*. Data shown is representative of at least 3 independent experiments with n=4 mice/group

## METHODS AND MATERIALS

### Animals

C57Bl/6J, B6;129S-*Tnf*^*tm1Gkl*^/J (*Tnfa*^*−/−*^), B6.129P2-*Nos2*^*tm1Lau*^/J (*Nos2*^*−/−*^), B6.129S2-*H2*^*dlAb1-Ea*^/J (*MHCII*^*−/−*^), B6.129S2-*Ighm*^*tm1Cgn*^/J (*μMT*^−/-^) mice were purchased from the Jackson Laboratories. Mice were maintained in specific pathogen free (SPF) conditions at the University of Toronto Division of Comparative Medicine animal facility. Mice were used at the age of 6-12 weeks. Experiments were conducted using age and sex matched littermate controls. Animals were housed in a closed caging system and provided with irradiated chow diet (Envigo Teklad 2918), non-acidified water (reverse-osmosis and UV-sterilized) with a 12hr light/dark cycle. Following colonization with *Tritrichomonas musculis* animals were housed in separated cages to avoid contamination with non-colonized mice. Animal experiments were approved by the Local Animal Care Committee (LACC) at the Faculty of Medicine, University of Toronto.

### Colonization of germ-free animals

Germ free C57Bl/6 mice were bred in isolators at the Microbiome Translational Center Gnotobiotic Facility at the Icahn School of Medicine. Mice 6-8 weeks of age were colonized by oral gavage with one of six strains of *B. ovatus* (cultured anaerobically in rich BHI media as in Yang et al.)(35) and then housed in barrier cages under aseptic conditions. Experiments were approved by the Institutional Animal Care and Use Committee in Icahn School of Medicine at Mount Sinai.

### *Tritrichomonas* Colonization

Purification of *T.mu* was performed as described in (5). The caecal content of *T.mu* colonized mice was harvested into sterile PBS and filtered through a 70μm cell strainer. Cecal contents were spun at 500g for 7min at 4°C. The supernatant was discarded, and the pellet was washed twice with sterile PBS. Protozoa were enriched through a percoll gradient and the interphase collected, washed and filtered through a 70μm cell strainer. The *T.mu* were sorted into sterile PBS on a BD Influx using the 100μm nozzle at 27psi at 4°C at a purity of >99%. Sorted *T.mu* were spun down and resuspended in sterile PBS prior to orally gavaged of 2×10^6^ cells into mice.

### Quantification of *T.mu* Colonization

Quantification of *T.mu* was perform as previously described (5). In brief, caeca were harvested, opened and the caecal content resuspended in 10ml of sterile PBS. Protozoa were counted using a hemocytometer.

### Isolation of Lymphocytes

Lamina propria (LP) lymphocytes were isolated as described in Burrows *et al*., 2019 (46). The large intestines were incubated in Hank’s balanced salt solution (HBSS w/o Ca^2+^Mg^2+^, GIBCO) with EDTA (5mM) and HEPES (5mM) for 10min at 37**°**C with mild agitation. Samples were vortexed for 10s and washed in 37**°**C HBSS buffer (HBSS with Ca^2+^Mg^2+^, GIBCO). Tissues were minced into ∼1mm pieces and transferred into 10mL of digestion buffer containing HBSS with Ca^2+^Mg^2+^, HEPES (5mM), FBS (2%), Collagenase and DNaseI. Digested samples were vortexed for 10s and filtered with a 70μm cell strainer and pellets, containing leukocytes resuspended and separated via percoll gradient centrifugation. The interphases were collected, washed once with media and spun down at 500g for 7min at 4°C prior to use in experiments.

Peyer’s patches (PPs) were rinsed in cold HBSS w/o Ca^2+^Mg^2+^, supplemented with EDTA (5mM) and HEPES (5mM) and incubated in the same solution for 10min at 37**°**C with mild agitation. Epithelial remains were removed, and PPs mashed through a 70μm cell strainer. Samples were washed once with FACS buffer (PBS, 5mM EDTA, 2% FBS) and spun down at 500g for 7min at 4°C prior to the use in experiments.

Mesenteric lymph nodes (MLNs) were dissected, and surrounding fat removed. MLNs were mashed through a 70μm cell strainer. Samples were washed once with FACS buffer and spun down at 500g for 7min at 4°C prior to the use in experiments.

Spleens were dissected and mashed through a 70μm cell strainer. Samples were washed once with PBS and spun down at 500g for 7min at 4°C. Supernatant was removed, and samples were incubated in 1mL of 1x red cell lysis (RCL) buffer for 3min at RT. Samples were washed once with FACS buffer and spun down at 500g for 7min at 4°C prior to the use in experiments.

Femur and tibia were collected and remaining soft tissues removed from the bones. Bone marrow was flushed out using cold PBS and samples washed once with PBS. Supernatant was removed, and samples were incubated in 1mL of 1x RCL buffer for 3min at RT. Samples were washed once with FACS buffer and spun down at 500g for 7min at 4°C prior to the use in experiments.

### Fecal Bacterial Processing

Mouse fecal samples were collected into 1mL of PBS in sterile Eppendorf tubes. Fecal weights were recorded. Samples were mechanical disrupted at 2500rpm for 3min using a bead disruptor (without beads). Samples rested on ice for 5min and were filtered using a 70μm filter. Filtered solutions were transferred into a new Eppendorf tube. Samples were spun at 50g for 3min and the supernatant transferred to a new Eppendorf tube. Samples were washed twice with PBS prior to use.

### Bacterial Flow Cytometry

Bacteria were resuspended in Bacteria Blocking Buffer (BBB: PBS, 20% (v/v) goat serum) and incubated at 4**°**C for 20 min. Samples were washed with Bacteria Staining Buffer (BSB: PBS, 1% BSA, 40 μM filtered) and spun at 8000g for 4min. Fecal bacterial samples from *Rag2*^*−/−*^ mice were incubated with 50μL of serum for 30min at 4**°**C and samples washed twice with BSB. Bacteria were stained with fluorophore-conjugated anti-IgA antibodies in BSB in the dark at 4**°**C for 30min, and washed twice with BSB at 8000g for 4min. Samples were further stained with PBS supplemented SytoBC (1:2000) at 4**°**C for 10min in the dark, washed twice, and resuspended in BSB for analysis.

### Flow Cytometry

*Surface Staining*: Cells were washed once with FACS buffer and spun down at 500g for 4min at 4°C. For CXCR5 surface staining, samples were incubated with anti-CXCR5 in FACS buffer (1/50 – RT – 45min). Samples were washed twice at 500g for 4min at 4°C, then incubated with corresponding surface antibody cocktails in FACS buffer at 4°C for 30min. Samples were washed twice and resuspended in FACS buffer for analysis. *Intracellular Staining*: Following surface staining, intracellular staining was performed using the eBioscience™ Foxp3 kit following the manufacturer’s recommendations. Samples were resuspended in intracellular staining buffer analysis. Flow cytometry was performed on a BD LSRFortessa™ X-20 and analyzed using FlowJo (v10.1).

### Fluorescent Activated Cell Sorting

*Lymphocyte FACS*: Lymphocyte sorting was performed on a BD FACSAria. Up to 5 million cells were sorted into 1mL of sorting buffer (SB – 50% FACS buffer - 50% FBS) for downstream applications. *Fecal Bacteria FACS*: Fecal bacteria sorting was performed on a BD FACSAria. Up to 2 million cells were sorted into 1mL of DNAzap (Invitrogen™) for downstream applications.

### Immunofluorescence Microscopy

*Colon Sample Preparation*: Following colon cleaning, samples were flushed with cold PBS. Colons were cut into four equal parts and placed into 2mL of fixative solution (2% PFA - 20% sucrose - 1X PBS). Samples were fixed for 1hr at RT and transferred into PBS on ice for 1hr. Samples were incubated in 40% sucrose/PBS overnight at 4°C. Samples were embedded into OCT and stored at −80°C. Tissues were sections at 7μm onto Fisherbrand™ Superfrost slides and slides stored at −20°C. *IF Staining Procedure*: Sections were brought to RT, circled with a PAP pen (Abcam) and air-dried for 15min. Sections were then rehydrated three times with PBS for 5 min and blocked with blocking buffer (10% BSA - 0.1% TritionX – PBS) for 1hr at RT in a humidified chamber. Sections were washed and stained with anti-IgA FITC (clone 11-44-2, Southern Biotech, 1:100) in blocking buffer in the dark for 30min at RT. Stained sections were washed and mounted with Fluoroshield mounting media with DAPI (Abcam). Slides were stored in the dark at 4°C or visualized immediately on a ZEISS Axio Imager.

### CD4 T cell Depletion

Mice were injected intra peritoneal (i.p.) with depleting anti-CD4 Ig 25mg/kg (clone GK1.5, BioXcell) in sterile PBS for three days prior to *T.mu* colonization and with 15mg/kg anti-CD4 Ig at D1, D7 and D14 post *T.mu* colonization.

### CD4^+^ T cell Transfer

Live, CD45^+^, CD3^+^, CD4^+^ splenic T cells were sorted on a BD FACSAria. Cells were resuspended in cold PBS and 2×10^5^ T cells were injected i.p. into age and sex-matched *Tcrb*^*−/−*^ mice. Two weeks post injection, mice were colonized with *T.mu*.

### ICOSL Blockade

Mice were injected i.p. with blocking anti-ICOSL Ig at 25mg/kg (HK5.3, BioXCell) one day prior to *T.mu* colonization. Post colonization, mice received 25mg/kg of anti-ICOSL Ig three times per week for the remainder of the experiments.

### Ovalbumin Treatment

Three weeks post *T.mu* colonization, mice were offer 1% OVA (Grade V, Millipore Sigma) in sterile drinking water *ad libidum*. OVA containing drinking water was freshly prepared and sterile filtered through a 20μm filter every two days for 3 weeks.

### Bone Marrow Chimeras

B cell-deficient *μMT*^−/-^ mice received a dose of 1300 cGy total body irradiation over two doses of 650 cGy, separated by a 12 hour period. Four hours following the last dose, recipient mice i.v. injected with a total of 5×10^6^ bone marrow leukocytes. Mice were treated with streptomycin-containing drinking water (1g/L) for the two weeks post bone marrow injections. Ten weeks after bone marrow injections, mice were colonized with *T.mu*.

### Total Ig ELISA

Mouse blood was collected isolated using Microvette CV 300 Z serum separation tubes (SARSTEDT Germany) according to the manufacturers protocol. 50 μL of anti-mouse Ig (SouthernBiotech – 1:2000) in PBS were added to 96-well high-bound Nunc MaxiSorp ELISA plates (BioLegend) and incubated at 4°C overnight. Plates were washed and blocked and serum and fecal samples (1:100, 1:10) were added. Samples were incubated for 1 hour at 37°C. Plates were washed and incubated with 50μL of anti-mouse IgA-HRP (1:1000) in assay buffer (SouthernBiotech). Plates were washed and developed using TMB substrate (BioLegend). Reaction was stopped using 50μL of H_2_SO_4_. Plates were read by spectrophotometry (450 nm excitation) and absorbance values were interpolated based on a purified IgA standard curve.

### Anti-OVA IgA ELISA

96-well ELISA plates (BioLegend) were coated with 50 μL of Ovalbumin from chicken egg white (Sigma-Aldrich, 1:1000 in 1x PBS) at 4°C overnight. Plates were washed and blocked prior to incubation with 100μl of serum (1:100 dilution) or fecal supernatant (1:10 dilution). Anti-OVA IgA standards (Chondrex) were run in parallel on each plate. Plates were then washed and incubated with 50μL of anti-mouse IgA-HRP (SouthernBiotech, 1:1000 in PBT), washed and developed using 50 μL of TMB substrate (BioLegend). Reaction was stopped using 50μl of H_2_SO_4_ and plates were read on a spectrophotometry (450 nm excitation) and absorbance values were interpolated based on a purified IgA standard curve.

### Anti-OVA and Total IgA ELISPOT

96-well filter plates (Millipore) were coated with 50 μL of Ovalbumin from chicken egg white (Sigma-Aldrich, 1:1000 in 1X PBS) and incubated at 4°C overnight. Plates were washed and blocked and 100μl of diluted cell samples in complete RPMI were added to the plates and titrated. Plates were incubated overnight at 37°C in a tissue culture chamber. The following day, plates were washed and 100μl of anti-mouse IgA-HRP (SouthernBiotech) added followed by incubation at 37°C for 2 hours. Plates were washed 2 times with PBST and incubated with mild shaking at RT for 10min during the third wash. Plates then were washed twice with PBST. 50 μl of AEC buffer (Vector laboratories; SK-4200, in distilled water) was added to each well, followed by distilled water to stop the reaction after 9 minutes. Plates were placed in the dark for drying overnight. The following day, red-coloured spots on each well’s membrane were counted as one ASC.

### DNA Isolation for Whole Fecal Samples

DNA isolation was performed using the QIAQuick qPCR purification kit (QIAGEN) with modifications. Fecal weights were measured, and flash-frozen pellets were defrosted on ice for 10min. SDS buffer (8g SDS – 30mL H_2_0 – 0.22*μm filtered*) and DNA buffer A (20mM Tris pH 8 – 2mM EDTA – 200mM NaCl) were prepared. SDS buffer and DNA buffer A were combined in a 1:1.41 ratio to prepare SDS-A buffer. Samples were transferred to cryogenic 2ml tubes and 400μ*l* zirconium beads, 550μl of phenol-chloroform, and 482μl of SDS-A buffer were mixed. Samples were bead beaten at 1800g for 5min. Samples were spun at 1800g for 5min, and the supernatant was transferred into a QIAGEN PCR purification plate containing 650μl of PM buffer (QIAQuick qPCR kit). Samples were washed twice with 900μl of PE buffer (QIAQuick qPCR kit). Samples were spun at 1800g for 5min to remove trace ethanol. To elute DNA, 100μl of DNAse/RNAse free H_2_O was added to each well. Samples were incubated at RT for 5min and spun at 1800g for 2min to collect DNA. DNA concentrations were quantified using the Qbit Quant-iT™ kit.

### RNA Isolation

Whole colonic tissue or purified *T.mu* were resuspended in 500μL of TRIzol™ and RNA isolated using the phenol/chloroform extraction. Recovered RNA was resuspended in 50μL of RNase-free water and stored at −80°C. For FACS sorted cells, RNA was purified in accordance to the RNeasy kit (QIAGEN). RNA concentration was quantified using a NanoDrop™.

### Reverse Transcription of mRNA

Total mRNA was converted to cDNA using the High-Capacity RNA-to-cDNA™ kit (Applied Biosystems). In brief, purified mRNA was mixed with 2X Reverse Transcription Buffer Mix, 20X Reverse Transcription Enzyme Mix and nuclease free H_2_O. The thermal cycling conditions were conducted as follows: 37*°*C for 60min - 95*°*C for 5min - infinite hold at 4*°*C. Controls for reverse transcription included a non-template control, no enzyme control and water-only control.

### Digital Droplet PCR (ddPCR)

ddPCR was performed following the QX200™ ddPCR™ EvaGreen Supermix kit instructions (Bio-Rad). In brief, 50ng of DNA template per reaction was mixed in nuclease-free ddPCR plates with 2X QX200™ ddPCR™ EvaGreen Supermix, 100nM of forward/reverse primer, 1μL of diluted restriction enzyme and nuclease-free water. Oil droplet generation was performed in a QX200 AutoDG Droplet Digital PCR System (Bio-Rad) using the QX200 Droplet Generation oil for EvaGreen (Bio-Rad, cat #1864005). The thermal cycling conditions were conducted as follows: Enzyme inactivation at 95*°*C for 5min (1 cycle) – denaturation at 95*°*C for 30s, annealing/extension at 60°C for 60s (40 cycles) – signal stabilization at for 4*°*C for 5min followed by 90*°*C for 5min (1 cycle) – hold at 4*°*C. Ramp rate between cycling steps was set at 2°C/sec. Samples were analyzed using the QX200 Droplet Reader (Bio-Rad) and analyzed using the QuantaSoft™ software (Bio-Rad). Values were reported as absolute concentration as copies/μL.

### Quantitative Real-Time PCR (qPCR)

qPCR was performed following the PowerTrack SYBR Green Master Mix™ kit instructions (Applied Biosystems). In brief, 10ng of DNA per reaction was mixed in nuclease-free qPCR plates with 2X PowerTrack Master Mix, 100nM of forward/reverse primer and nuclease-free water. The thermal cycling conditions were conducted as follows: Enzyme inactivation at 95*°*C for 2min (1 cycle) – denaturation at 95*°*C for 15s, annealing/extension at 60*°*C for 60s (40 cycles). Controls for qPCR included a non-template control, no master mix control and water-only control. Relative expression was calculated using the delta-delta Ct method.

### 16S rDNA Analysis

Purified DNA from mouse fecal pellets or sorted cells was subjected to 16S variable region 4 PCR amplification using barcoded 515F and 806R primers (47) and the KAPA2G Robust HotStart ReadyMix (KAPA Biosystems), with the following cycling conditions: 95**°**C for 3 min, 24x cycles of 95**°**C for 15 sec, 50**°**C for 15 sec and 72**°**C for 15 sec, followed by a 5 min 72**°**C extension. All reactions were performed in triplicate and pooled. The amplicon libraries were combined and purified using Ampure XP beads, and sequenced at the Center for the Analysis of Genome Evolution and Function (CAGEF) at the University of Toronto on an Illumina MiSeq using V2 (150bp x2) chemistry, according to manufacturer instructions (Illumina, San Diego, CA). Amplicon sequences were quality filtered and classified using the UNOISE pipeline, available through USEARCH v11.0.667 and vsearch v2.10.4 (48–51). Briefly, low quality bases were trimmed using cutadapt v.1.18 and read pairs were merged using vsearch with assemble lengths set between 243 and 263. Amplicons were denoised and filtered for chimeras, and clustered to 99% identity OTUs. Taxonomies were assigned using SINTAX and the Ribosomal Database Project (RDP) database version 16, with a minimum confidence cutoff of 0.8 (52). A phylogenetic tree was generated using FastTree (53).

Compositional and diversity analyses of bacterial data were carried out using R-packages Phyloseq 1.26.1 and vegan v.2.5. Alpha diversity metrics (observed OTUs and the Shannon Diversity Index) were determined from averages of 100 independent datasets rarefied to the minimum sample read depth (19,427), and group differences were evaluated using the generalized linear models (glm) function in R. Beta diversities (compositional differences among samples) were calculated based on Bray-Curtis dissimilarity scores using rarefied OTU data (filtered for OTUs with minimum 5 reads), and tested for group differences using the adonis function in vegan. Differential taxon abundance among groups was determined with DESeq2 v.1.22.2. Heatmaps of bacterial abundance were generated using variance stabilized counts with the package pheatmap v1.0.12.

### Statistical Analysis

Statistics were performed using Prism 7 software. Error bars are presented using the standard of error of the mean (SEM). Student’s t-test, Mann-Whitney, one-way ANOVA and two-way ANOVA tests using Prism 7 were used to determine statistical significance where appropriate. Significance was indicated as follows: * = p < 0.05, ** = p < 0.01, *** = p < 0.001, **** = p < 0.0001, ns = not significant.

### Antibodies

| Fluorophore | Antibody | Clone | Company | Reference # |
| --- | --- | --- | --- | --- |
| APC | Rat Anti-Mouse B220 (CD45R) | RA3-6B2 | BioLegend | 103212 |
| APC | Rat Anti-Mouse PD-1 (CD279) | 29F.1A12 | BioLegend | 135210 |
| AF647 | Goat Anti-Mouse IgA | N/A | Southern Biotech | 1040-31 |
| eF450 | Rat Anti-Mouse CD3 | 17A2 | eBioscience | 48-0032-82 |
| eF450 | Rat Anti-Mouse MHC-II (I-A/I-E) | M5/114.15.2 | BioLegend | 107607 |
| FITC | Rat Anti-Mouse CD40L (CD154) | SA047C3 | BioLegend | 157005 |
| FITC | Rat Anti-Mouse CD4 | GK1.5 | BioLegend | 100406 |
| FITC | Rat Anti-Mouse CD40L (CD154) | SA047C3 | BioLegend | 157006 |
| FITC | Anti-Mouse FAS (CD95) | SA367H8 | BioLegend | 152606 |
| FITC | Goat Anti-Mouse IgA | N/A | Southern Biotech | 1165-02 |
| FITC | Rat Anti-Mouse Ki-67 | 16A8 | BioLegend | 652410 |
| FITC | Rat Anti-Mouse/Rat XCR1 | ZET | BioLegend | 148210 |
| FITC | SYTOBC Green Fluorescent<br>Nucleic Acid Stain | N/A | ThermoFisher | S34855 |
| APC | Rat Anti-Mouse B220 | RA3-6B2 | BioLegend | 103231 |
| APC | Rat Anti-Mouse PD-1 | RMP1-30 | BioLegend | 109111 |
| APC | Rat Anti-Mouse CD40 | 3/23 | BioLegend | 124612 |
| APC Cy7 | Anti-Mouse CD45.2 | 104 | BioLegend | 109824 |
| PE | Rat Anti-Mouse CD197 (CCR7) | 4B12 | BioLegend | 120106 |
| PE | Rat Anti-Mouse CD138 | 281-2 | BioLegend | 142504 |
| PE | Hamster Anti-Mouse CD154 | MR1 | BioLegend | 106505 |
| PE | Rat Anti-Mouse GL-7 | GL7 | BioLegend | 144608 |
| PE | Rat Anti-Mouse Foxp3 | MF-14 | BioLegend | 126404 |
| PE | Rat Anti-Mouse CCR7 | 4B12 | BioLegend | 120106 |
| PE | Rat Anti-Mouse ICOSL | HK5.3 | BioLegend | 107405 |
| PE Cy7 | Anti-Mouse/Human CD11b | M1/70 | BioLegend | 101216 |
| PE Cy7 | Hamster Anti-mouse ICOS | C398.4A | Biolegend | 313520 |
| PE Cy7 | Rat Anti-Mouse CD4 | GK1.5 | BioLegend | 100422 |
| PerCP Cy5.5 | Rat Anti-Mouse IgM | RMM-1 | BioLegend | 406512 |
| PerCP e710 | Rat Anti-Mouse CXCR5 | SPRCL5 | eBioscience | 46-7185-82 |
| BV 605 | Rat Anti-Mouse B220 | RA3-6B2 | BioLegend | 103243 |
| eFluor™ 506 | Fixable Viability | N/A | eBioscience | 65-0866-14 |
| HRP | Goat Anti-Mouse IgA | N/A | Southern Biotech | 1040-05 |
| N/A | InVivoMAb anti-mouse ICOSL | HK5.3, | BioXCell | BE0028 |
| N/A | InVivoMAb anti-mouse CD4 | GK1.5 | BioXCell | BE0003-1 |

### Reagents and Resources

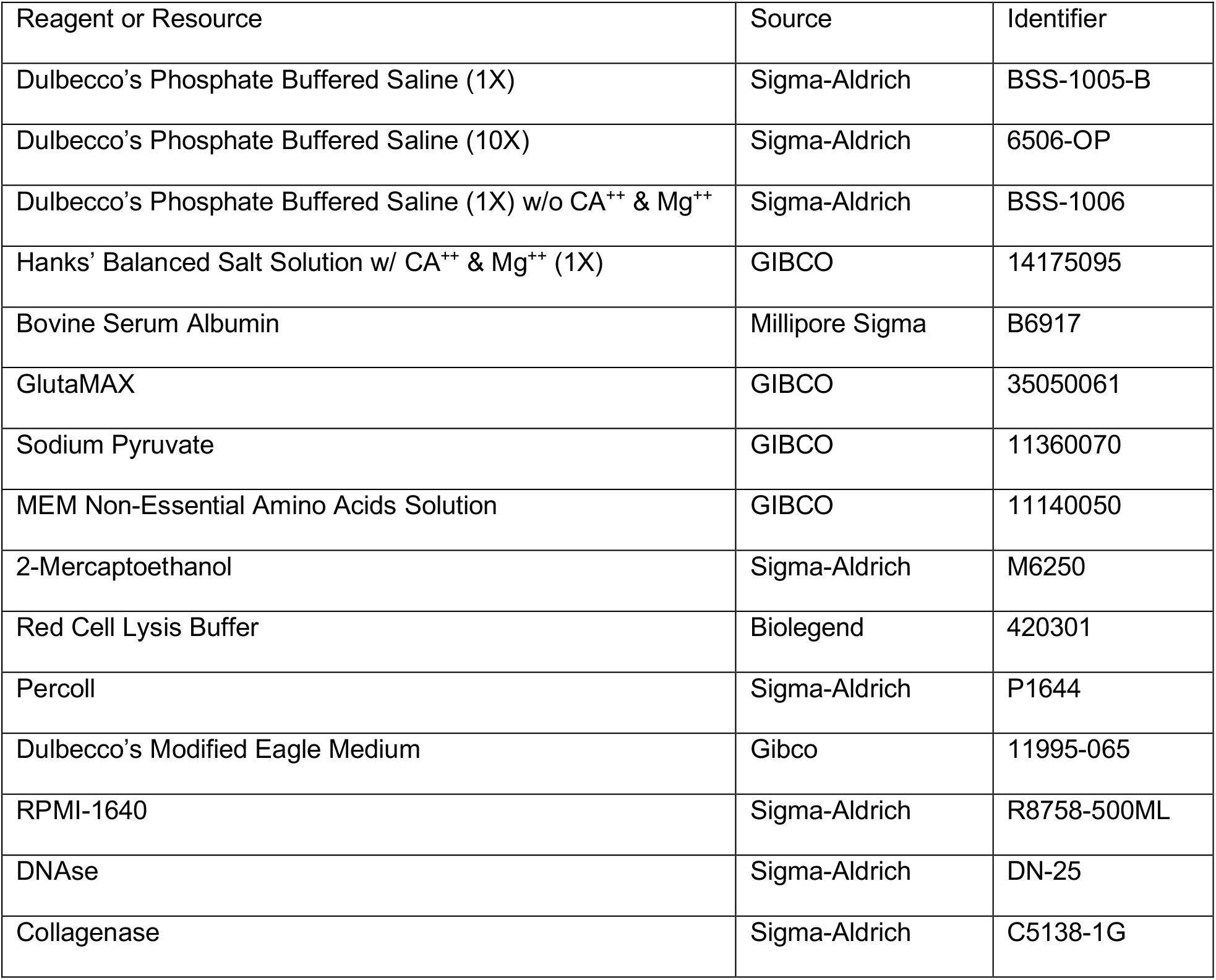
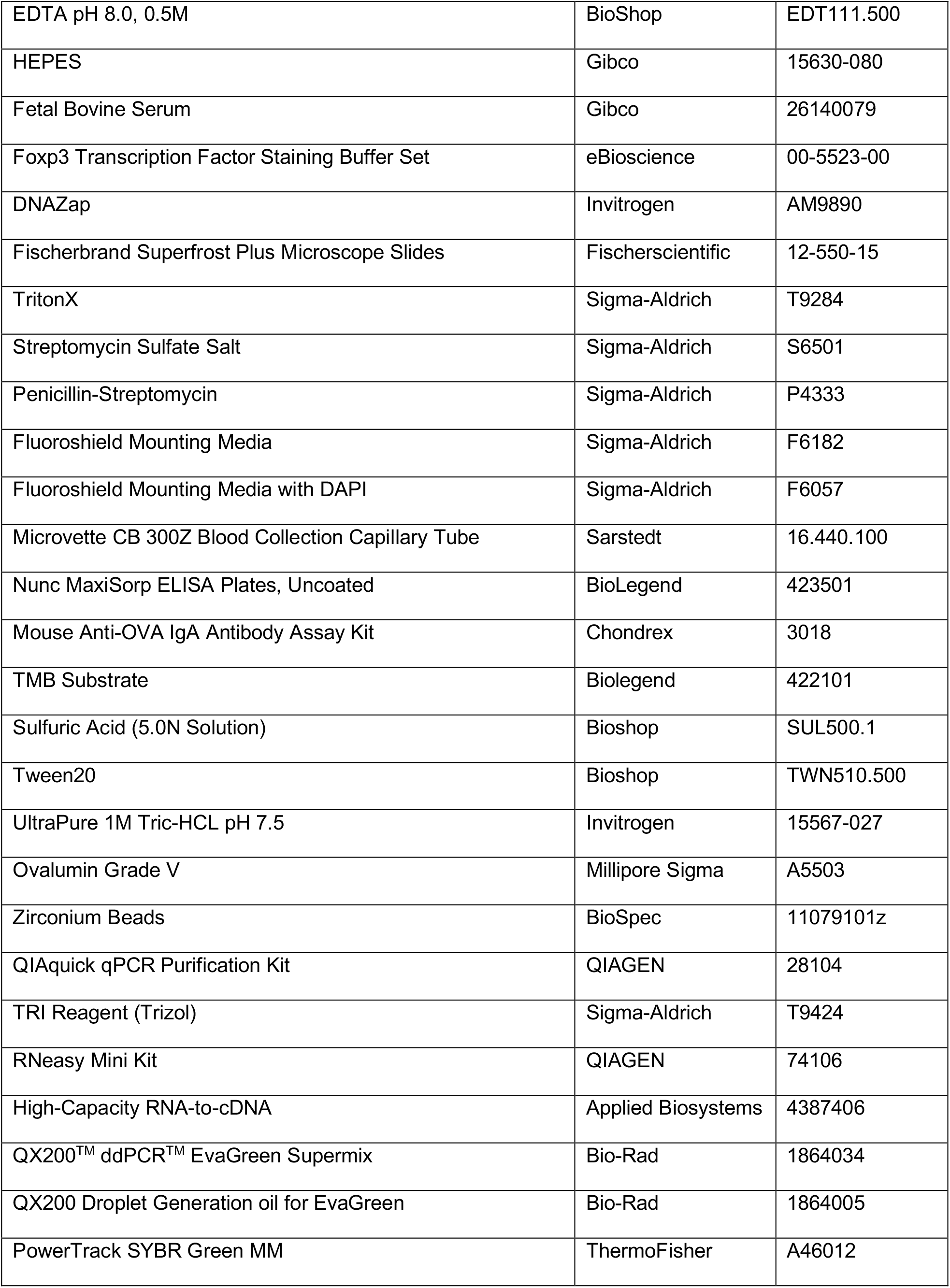
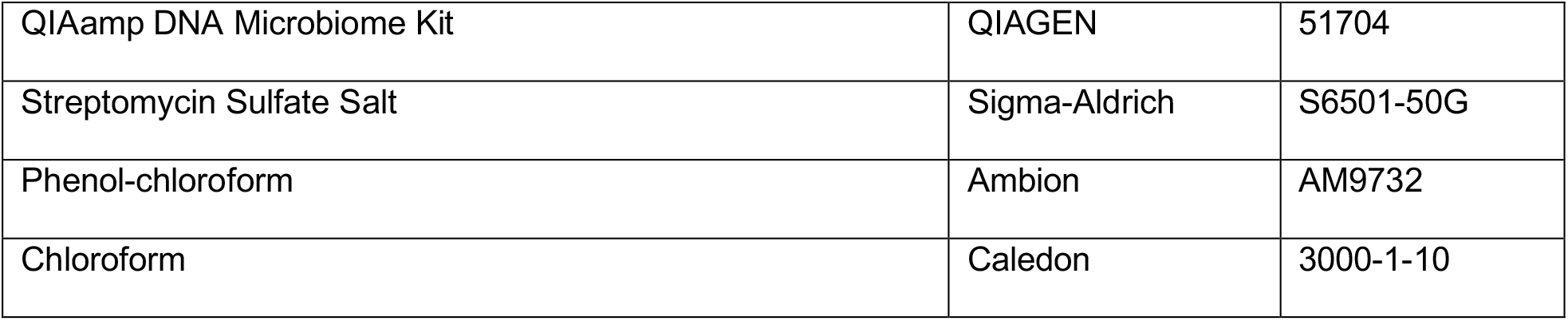

| Reagent or Resource | Source | Identifier |
| --- | --- | --- |
| Dulbecco's Phosphate Buffered Saline (1X) | Sigma-Aldrich | BSS-1005-B |
| Dulbecco's Phosphate Buffered Saline (10X) | Sigma-Aldrich | 6506-OP |
| Dulbecco's Phosphate Buffered Saline (1X) w/o $Ca^{++}$ & $Mg^{++}$ | Sigma-Aldrich | BSS-1006 |
| Hanks' Balanced Salt Solution w/ $Ca^{++}$ & $Mg^{++}$ (1X) | GIBCO | 14175095 |
| Bovine Serum Albumin | Millipore Sigma | B6917 |
| GlutaMAX | GIBCO | 35050061 |
| Sodium Pyruvate | GIBCO | 11360070 |
| MEM Non-Essential Amino Acids Solution | GIBCO | 11140050 |
| 2-Mercaptoethanol | Sigma-Aldrich | M6250 |
| Red Cell Lysis Buffer | Biolegend | 420301 |
| Percoll | Sigma-Aldrich | P1644 |
| Dulbecco's Modified Eagle Medium | Gibco | 11995-065 |
| RPMI-1640 | Sigma-Aldrich | R8758-500ML |
| DNase | Sigma-Aldrich | DN-25 |
| Collagenase | Sigma-Aldrich | C5138-1G |
| EDTA pH 8.0, 0.5M | BioShop | EDT111.500 |
| HEPES | Gibco | 15630-080 |
| Fetal Bovine Serum | Gibco | 26140079 |
| Foxp3 Transcription Factor Staining Buffer Set | eBioscience | 00-5523-00 |
| DNAZap | Invitrogen | AM9890 |
| Fischerbrand Superfrost Plus Microscope Slides | Fischerscientific | 12-550-15 |
| TritonX | Sigma-Aldrich | T9284 |
| Streptomycin Sulfate Salt | Sigma-Aldrich | S6501 |
| Penicillin-Streptomycin | Sigma-Aldrich | P4333 |
| Fluoroshield Mounting Media | Sigma-Aldrich | F6182 |
| Fluoroshield Mounting Media with DAPI | Sigma-Aldrich | F6057 |
| Microvette CB 300Z Blood Collection Capillary Tube | Sarstedt | 16.440.100 |
| Nunc MaxiSorp ELISA Plates, Uncoated | BioLegend | 423501 |
| Mouse Anti-OVA IgA Antibody Assay Kit | Chondrex | 3018 |
| TMB Substrate | Biolegend | 422101 |
| Sulfuric Acid (5.0N Solution) | Bioshop | SUL500.1 |
| Tween20 | Bioshop | TWN510.500 |
| UltraPure 1M Tris-HCL pH 7.5 | Invitrogen | 15567-027 |
| Ovalumin Grade V | Millipore Sigma | A5503 |
| Zirconium Beads | BioSpec | 11079101z |
| QIAquick qPCR Purification Kit | QIAGEN | 28104 |
| TRI Reagent (Trizol) | Sigma-Aldrich | T9424 |
| RNeasy Mini Kit | QIAGEN | 74106 |
| High-Capacity RNA-to-cDNA | Applied Biosystems | 4387406 |
| QX200™ ddPCR™ EvaGreen Supermix | Bio-Rad | 1864034 |
| QX200 Droplet Generation oil for EvaGreen | Bio-Rad | 1864005 |
| PowerTrack SYBR Green MM | ThermoFisher | A46012 |
| QIAamp DNA Microbiome Kit | QIAGEN | 51704 |
| Streptomycin Sulfate Salt | Sigma-Aldrich | S6501-50G |
| Phenol-chloroform | Ambion | AM9732 |
| Chloroform | Caledon | 3000-1-10 |

### Primers

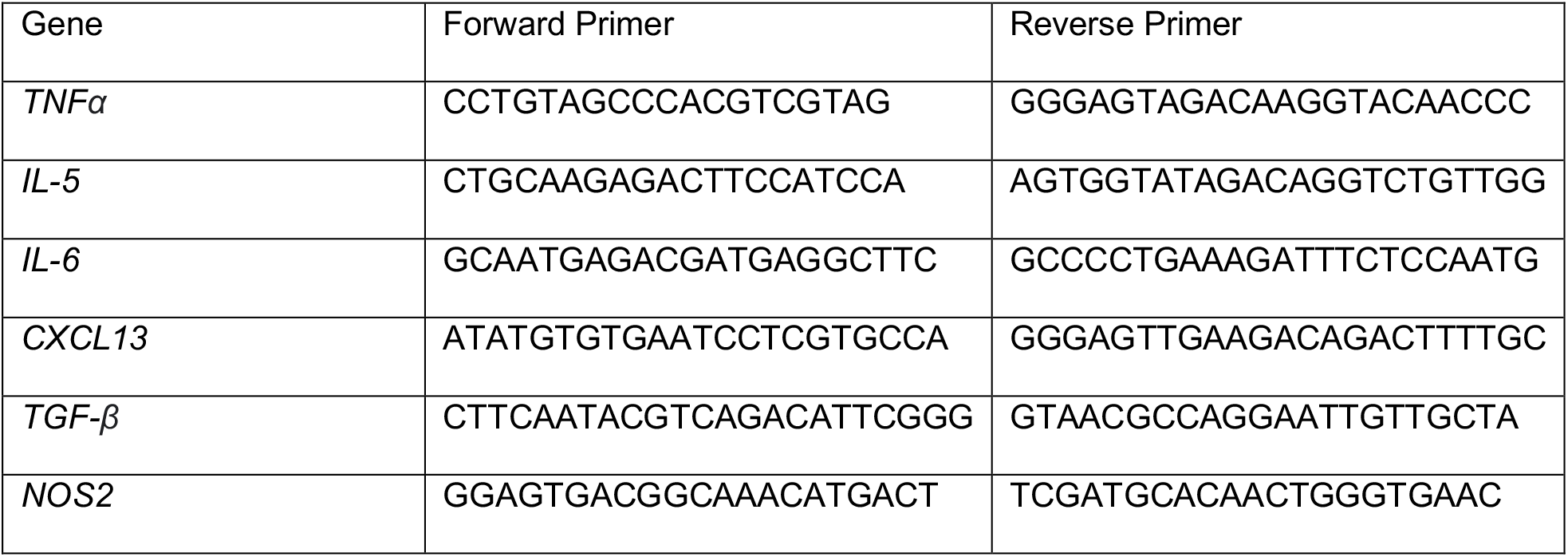

## References

1. Blander JM, Longman RS, Iliev ID, Sonnenberg GF, Artis D. Regulation of inflammation by microbiota interactions with the host. Nat Immunol. 2017;18(8):851–60.

2. Cao EY, Mortha A, editors. Tritrichomonas spp. and Their Impact on Gut Immune Homeostasis. Eukaryome Impact on Human Intestine Homeostasis and Mucosal Immunology; 2020 2020//; Cham: Springer International Publishing.

3. Howitt MR, Lavoie S, Michaud M, Blum AM, Tran SV, Weinstock JV, et al. Tuft cells, taste-chemosensory cells, orchestrate parasite type 2 immunity in the gut. Science. 2016;351(6279):1329–33.

4. Nadjsombati MS, McGinty JW, Lyons-Cohen MR, Jaffe JB, DiPeso L, Schneider C, et al. Detection of Succinate by Intestinal Tuft Cells Triggers a Type 2 Innate Immune Circuit. Immunity. 2018;49(1):33–41 e7.

5. Chudnovskiy A, Mortha A, Kana V, Kennard A, Ramirez JD, Rahman A, et al. Host-Protozoan Interactions Protect from Mucosal Infections through Activation of the Inflammasome. Cell. 2016;167(2):444–56 e14.

6. Escalante NK, Lemire P, Cruz Tleugabulova M, Prescott D, Mortha A, Streutker CJ, et al. The common mouse protozoa Tritrichomonas muris alters mucosal T cell homeostasis and colitis susceptibility. J Exp Med. 2016;213(13):2841–50.

7. Chiaranunt P, Burrows K, Ngai L, Cao EY, Liang H, Tai SL, et al. NLRP1B and NLRP3 Control the Host Response following Colonization with the Commensal Protist Tritrichomonas musculis. J Immunol. 2022;208(7):1782–9.

8. Schneider C, O’Leary CE, von Moltke J, Liang HE, Ang QY, Turnbaugh PJ, et al. A Metabolite-Triggered Tuft Cell-ILC2 Circuit Drives Small Intestinal Remodeling. Cell. 2018;174(2):271–84 e14.

9. Rojas OL, Probstel AK, Porfilio EA, Wang AA, Charabati M, Sun T, et al. Recirculating Intestinal IgA-Producing Cells Regulate Neuroinflammation via IL-10. Cell. 2019;176(3):610–24 e18.

10. Belkaid Hand, Y TW. Role of the microbiota in immunity and inflammation. Cell. 2014;157(1):121–41.

11. Cerutti A, Chen K, Chorny A. Immunoglobulin responses at the mucosal interface. Annu Rev Immunol. 2011;29:273–93.

12. Johansen FE, Kaetzel CS. Regulation of the polymeric immunoglobulin receptor and IgA transport: new advances in environmental factors that stimulate pIgR expression and its role in mucosal immunity. Mucosal Immunol. 2011;4(6):598–602.

13. Brandtzaeg P, Prydz H. Direct evidence for an integrated function of J chain and secretory component in epithelial transport of immunoglobulins. Nature. 1984;311(5981):71–3.

14. Brandtzaeg P. Mucosal immunity: induction, dissemination, and effector functions. Scand J Immunol. 2009;70(6):505–15.

15. Isho B, Florescu A, Wang AA, Gommerman JL. Fantastic IgA plasma cells and where to find them. Immunol Rev. 2021;303(1):119–37.

16. Fadlallah J, Sterlin D, Fieschi C, Parizot C, Dorgham K, El Kafsi H, et al. Synergistic convergence of microbiota-specific systemic IgG and secretory IgA. J Allergy Clin Immunol. 2019;143(4):1575–85 e4.

17. Cerutti A. The regulation of IgA class switching. Nat Rev Immunol. 2008;8(6):421–34.

18. Li X, Zhang YK, Yin B, Liang JB, Jiang F, Wu WX. Toll-Like Receptor 2 (TLR2) and TLR4 Mediate the IgA Immune Response Induced by Mycoplasma hyopneumoniae. Infect Immun. 2019;88(1).

19. Fagarasan S, Kawamoto S, Kanagawa O, Suzuki K. Adaptive immune regulation in the gut: T cell-dependent and T cell-independent IgA synthesis. Annu Rev Immunol. 2010;28:243–73.

20. Macpherson AJ, Uhr T. Induction of protective IgA by intestinal dendritic cells carrying commensal bacteria. Science. 2004;303(5664):1662–5.

21. Tezuka H, Ohteki T. Regulation of IgA Production by Intestinal Dendritic Cells and Related Cells. Front Immunol. 2019;10:1891.

22. Hou S, Landego I, Jayachandran N, Miller A, Gibson IW, Ambrose C, et al. Follicular dendritic cell secreted protein FDC-SP controls IgA production. Mucosal Immunol. 2014;7(4):948–57.

23. Bunker JJ, Flynn TM, Koval JC, Shaw DG, Meisel M, McDonald BD, et al. Innate and Adaptive Humoral Responses Coat Distinct Commensal Bacteria with Immunoglobulin A. Immunity. 2015;43(3):541–53.

24. Li H, Limenitakis JP, Greiff V, Yilmaz B, Scharen O, Urbaniak C, et al. Mucosal or systemic microbiota exposures shape the B cell repertoire. Nature. 2020;584(7820):274–8.

25. Hapfelmeier S, Lawson MA, Slack E, Kirundi JK, Stoel M, Heikenwalder M, et al. Reversible microbial colonization of germ-free mice reveals the dynamics of IgA immune responses. Science. 2010;328(5986):1705–9.

26. Palm NW, de Zoete MR, Cullen TW, Barry NA, Stefanowski J, Hao L, et al. Immunoglobulin A coating identifies colitogenic bacteria in inflammatory bowel disease. Cell. 2014;158(5):1000–10.

27. Sanderson RD, Lalor P, Bernfield M. B lymphocytes express and lose syndecan at specific stages of differentiation. Cell Regul. 1989;1(1):27–35.

28. Pabst O. New concepts in the generation and functions of IgA. Nat Rev Immunol. 2012;12(12):821–32.

29. Cazac BB, Roes J. TGF-beta receptor controls B cell responsiveness and induction of IgA in vivo. Immunity. 2000;13(4):443–51.

30. Wan YY, Flavell RA. Regulatory T-cell functions are subverted and converted owing to attenuated Foxp3 expression. Nature. 2007;445(7129):766–70.

31. Tezuka H, Abe Y, Iwata M, Takeuchi H, Ishikawa H, Matsushita M, et al. Regulation of IgA production by naturally occurring TNF/iNOS-producing dendritic cells. Nature. 2007;448(7156):929–33.

32. Muramatsu M, Kinoshita K, Fagarasan S, Yamada S, Shinkai Y, Honjo T. Class switch recombination and hypermutation require activation-induced cytidine deaminase (AID), a potential RNA editing enzyme. Cell. 2000;102(5):553–63.

33. Odegard JM, Marks BR, DiPlacido LD, Poholek AC, Kono DH, Dong C, et al. ICOS-dependent extrafollicular helper T cells elicit IgG production via IL-21 in systemic autoimmunity. J Exp Med. 2008;205(12):2873–86.

34. Landuyt AE, Klocke BJ, Duck LW, Kemp KM, Muir RQ, Jennings MS, et al. ICOS ligand and IL-10 synergize to promote host-microbiota mutualism. Proc Natl Acad Sci U S A. 2021;118(13).

35. Yang C, Mogno I, Contijoch EJ, Borgerding JN, Aggarwala V, Li Z, et al. Fecal IgA Levels Are Determined by Strain-Level Differences in Bacteroides ovatus and Are Modifiable by Gut Microbiota Manipulation. Cell Host Microbe. 2020;27(3):467–75 e6.

36. Neutra MR, Pringault E, Kraehenbuhl JP. Antigen sampling across epithelial barriers and induction of mucosal immune responses. Annu Rev Immunol. 1996;14:275–300.

37. Ise W, Fujii K, Shiroguchi K, Ito A, Kometani K, Takeda K, et al. T Follicular Helper Cell-Germinal Center B Cell Interaction Strength Regulates Entry into Plasma Cell or Recycling Germinal Center Cell Fate. Immunity. 2018;48(4):702–15 e4.

38. Deenick EK, Chan A, Ma CS, Gatto D, Schwartzberg PL, Brink R, et al. Follicular helper T cell differentiation requires continuous antigen presentation that is independent of unique B cell signaling. Immunity. 2010;33(2):241–53.

39. Thouvenel CD, Fontana MF, Netland J, Krishnamurty AT, Takehara KK, Chen Y, et al. Multimeric antibodies from antigen-specific human IgM+ memory B cells restrict Plasmodium parasites. J Exp Med. 2021;218(4).

40. Mei HE, Yoshida T, Sime W, Hiepe F, Thiele K, Manz RA, et al. Blood-borne human plasma cells in steady state are derived from mucosal immune responses. Blood. 2009;113(11):2461–9.

41. Israf DA, Lajis NH, Somchit MN, Sulaiman MR. Enhancement of ovalbumin-specific IgA responses via oral boosting with antigen co-administered with an aqueous Solanum torvum extract. Life Sci. 2004;75(4):397–406.

42. Tan J, Cho H, Pholcharee T, Pereira LS, Doumbo S, Doumtabe D, et al. Functional human IgA targets a conserved site on malaria sporozoites. Sci Transl Med. 2021;13(599).

43. Reperant JM, Naciri M, Chardes T, Bout DT. Immunological characterization of a 17-kDa antigen from Cryptosporidium parvum recognized early by mucosal IgA antibodies. FEMS Microbiol Lett. 1992;78(1):7–14.

44. Ma CS, Deenick EK, Batten M, Tangye SG. The origins, function, and regulation of T follicular helper cells. J Exp Med. 2012;209(7):1241–53.

45. Boyaka PN. Inducing Mucosal IgA: A Challenge for Vaccine Adjuvants and Delivery Systems. J Immunol. 2017;199(1):9–16.

46. Burrows K, Chiaranunt P, Ngai L, Mortha A. Rapid isolation of mouse ILCs from murine intestinal tissues. Methods Enzymol. 2020;631:305–27.

47. Caporaso JG, Lauber CL, Walters WA, Berg-Lyons D, Huntley J, Fierer N, et al. Ultra-high-throughput microbial community analysis on the Illumina HiSeq and MiSeq platforms. ISME J. 2012;6(8):1621–4.

48. Edgar RC, Flyvbjerg H. Error filtering, pair assembly and error correction for next-generation sequencing reads. Bioinformatics. 2015;31(21):3476–82.

49. Edgar RC. UPARSE: highly accurate OTU sequences from microbial amplicon reads. Nat Methods. 2013;10(10):996–8.

50. Edgar RC. Search and clustering orders of magnitude faster than BLAST. Bioinformatics. 2010;26(19):2460–1.

51. Rognes T, Flouri T, Nichols B, Quince C, Mahe F. VSEARCH: a versatile open source tool for metagenomics. PeerJ. 2016;4:e2584.

52. Wang Q, Garrity GM, Tiedje JM, Cole JR. Naive Bayesian classifier for rapid assignment of rRNA sequences into the new bacterial taxonomy. Appl Environ Microbiol. 2007;73(16):5261–7.

53. Price MN, Dehal PS, Arkin AP. FastTree: computing large minimum evolution trees with profiles instead of a distance matrix. Mol Biol Evol. 2009;26(7):1641–50.

