## Supplemental Figures for "The protozoan commensal *Tritrichomonas musculis* is a natural adjuvant for mucosal IgA"

### supplementary Figure 1. Unaltered Tfh cell phenotype in the presence of *T.mu*

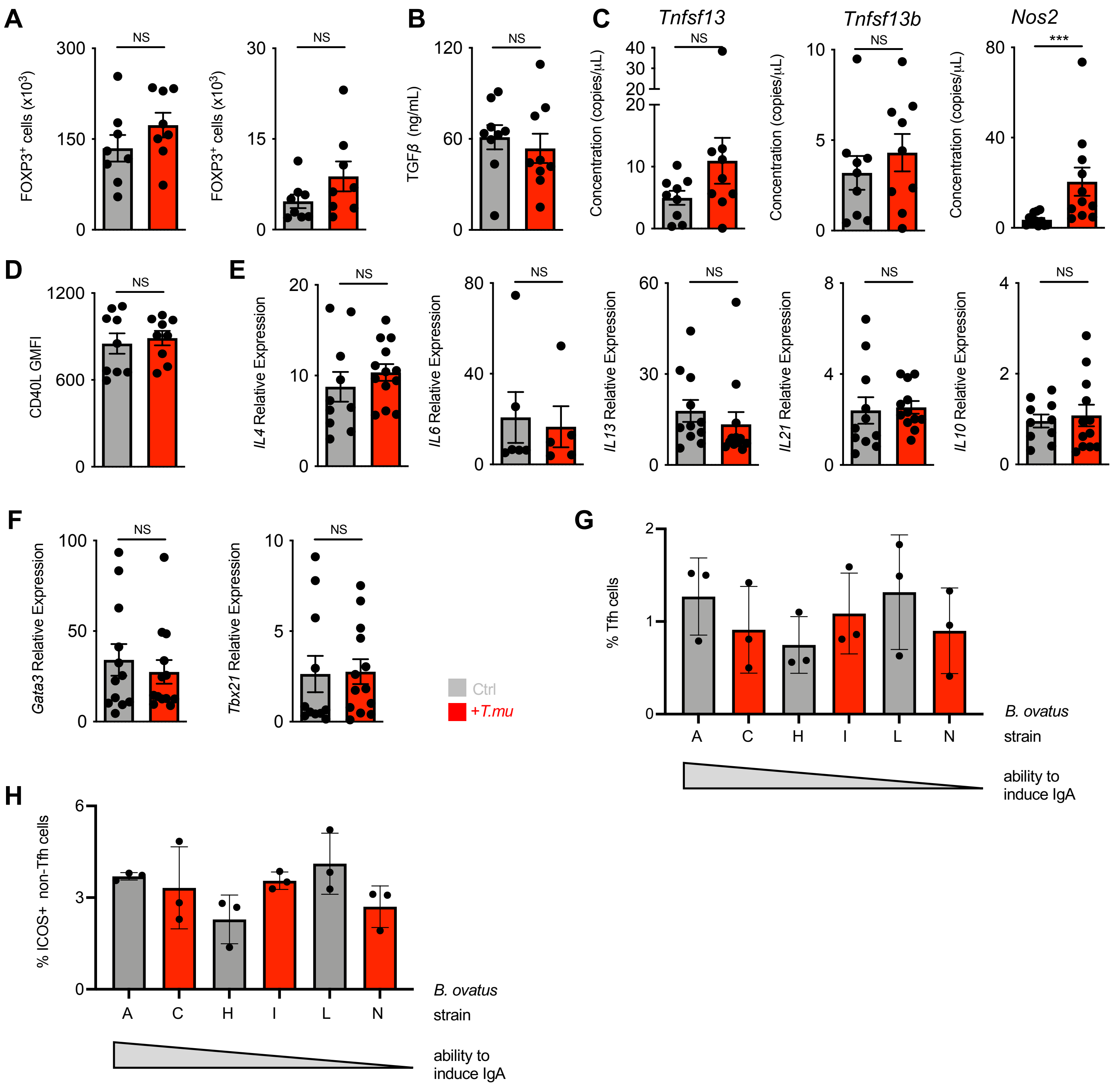

**supplementary Figure 2. Anticipatory shift in anti-bacterial IgA reactivity following colonization with *T.mu.***

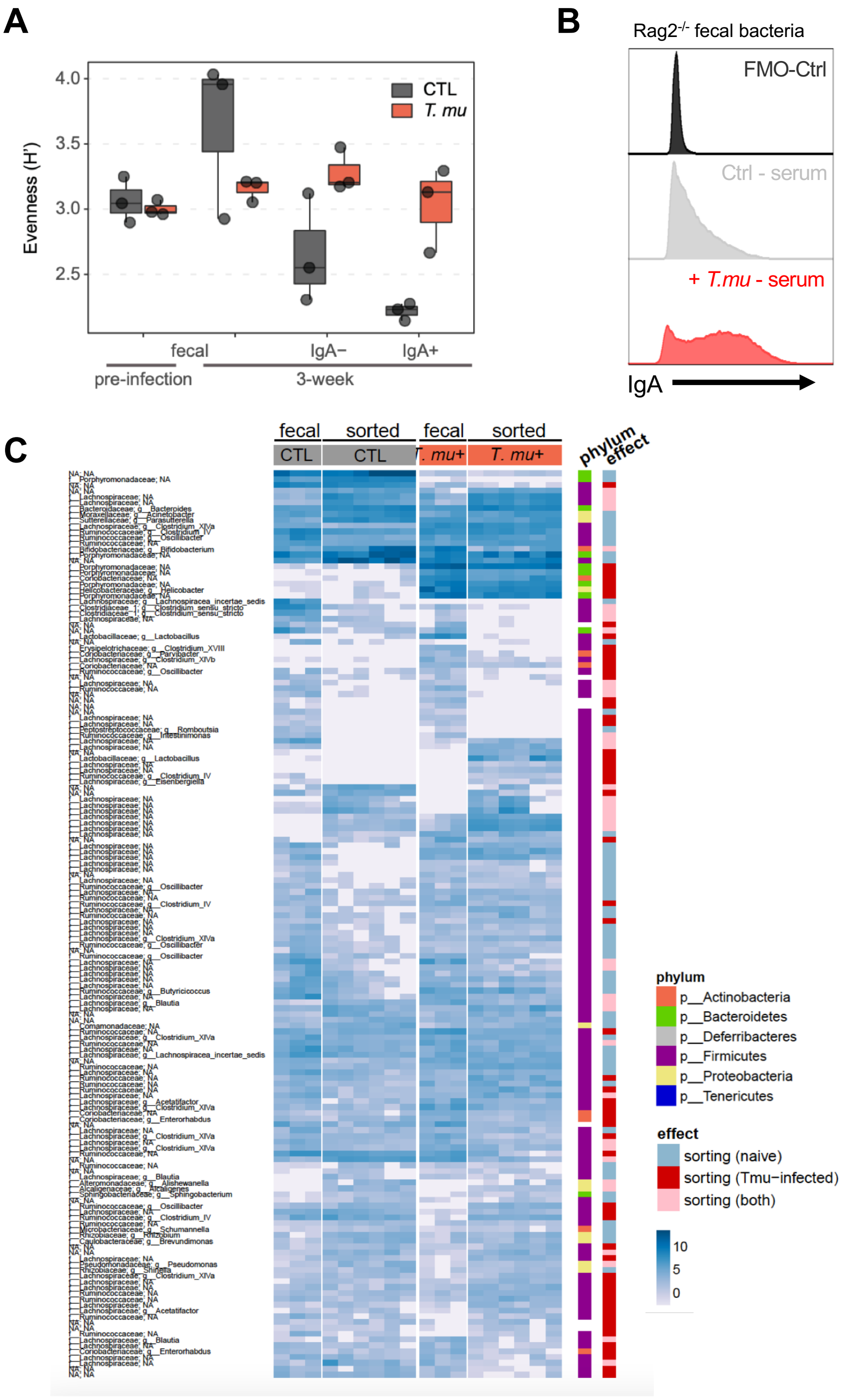

**supplementary Figure 3. B cell-dependent Tfh response in *T.mu*-colonized mice.**

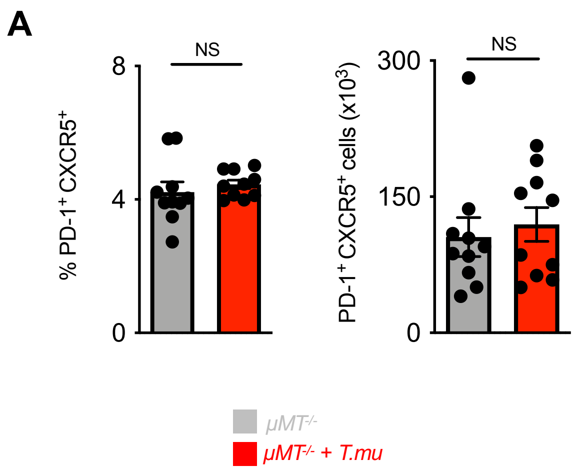

**supplementary Figure 4. Exposure to oral Ovalbumin does not alter the *T.mu*-driven IgA levels.**

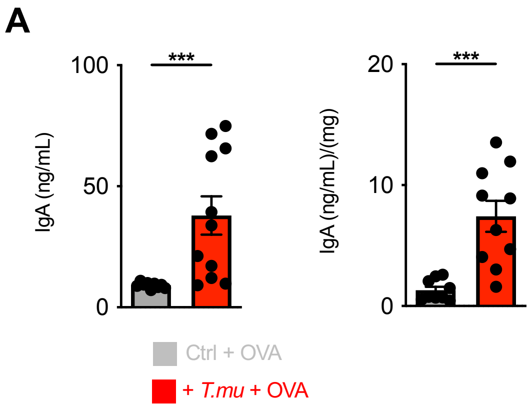
